# Whole Genome Assembly and Annotation of Northern Wild Rice, *Zizania palustris* L., Supports a Whole Genome Duplication in the *Zizania* Genus

**DOI:** 10.1101/2021.03.12.435103

**Authors:** Matthew Haas, Thomas Kono, Marissa Macchietto, Reneth Millas, Lillian McGilp, Mingqin Shao, Jacques Duquette, Candice N. Hirsch, Jennifer Kimball

## Abstract

Northern Wild Rice (NWR; *Zizania palustris* L.) is an aquatic grass native to North America that is notable for its nutritious grain. This is an important species with ecological, cultural, and agricultural significance, specifically in the Great Lakes region of the United States. Using long- and short-range sequencing, Hi-C scaffolding, and RNA-seq data from eight tissues, we generated an annotated whole genome *de novo* assembly of NWR. The assembly is 1.29 Gb, highly repetitive (∼76.0%), and contains 46,421 putative protein-coding genes. The expansion of retrotransposons within the genome and a whole genome duplication prior to the *Zizania-Oryza* speciation event have both led to an increase in genome size of NWR in comparison with *O. sativa* and *Z. latifolia*. Both events depict a genome rapidly undergoing change over a short evolutionary time. Comparative analyses revealed conservation of large syntenic blocks with *Oryza sativa* L., which were used to identify putative seed shattering genes. Estimates of divergence times revealed the *Zizania* genus diverged from *Oryza* ∼26-30 million years ago (MYA), while NWR and *Zizania latifolia* diverged from one another ∼6-8 MYA. Comparative genomics confirmed evidence of a whole genome duplication in the *Zizania* genus and provided support that the event was prior to the NWR-*Z. latifolia* speciation event. This high-quality genome assembly and annotation provides a valuable resource for comparative genomics in the Oryzeae tribe and provides an important resource for future conservation and breeding efforts of NWR.

## INTRODUCTION

Northern Wild Rice (NWR; *Zizania palustris* L.) is a diploid (2*n=*2*x=*30), annual, aquatic grass endemic to the Eastern Temperate and Northern Forest ecoregions of North America. NWR is a species with ecological, cultural, and agricultural significance, particularly in the Great Lakes region of the United States and Canada. In its native habitat, it is a vital component of aquatic ecosystems, providing food and shelter for a variety of species (Chambliss, 1940; Rogosin, 1954; Fannucchi, 1983). However, the species faces serious challenges due to habitat destruction, hydrological changes, and climate change (Pillsbury and McGuire, 2009; Drewes and Silbernagel, 2012). Also known as Manoomin or Psiŋ, NWR is a sacred food of Indigenous peoples living in the Great Lakes region, who harvest the grain for use in their daily lives and ceremonies, as barter in their trade economy, and for commercial sales (Andow *et al*., 2009). NWR is also considered a high-value specialty crop that is commercially cultivated in irrigated paddies, predominantly in Minnesota and California. It is prized for its nutritious grain, which has 2× the protein, 5× the dietary fiber, and ∼2× the essential amino acid content of white rice, *Oryza sativa* L. (Terrell and Wiser, 1975; Zhai *et al*., 1994; Surendiran *et al*., 2014).

As calls for improved conservation strategies of declining natural stands rise and commercial growers continue to face agronomic challenges, there is a growing need to expand the species’ genomic resources. In particular, NWR harbors several unique characteristics that pose challenges to both conservation and breeding schemes. The species’ seeds, for example, are intermediately recalcitrant or desiccation intolerant, which limits seed viability in *ex-situ* storage to 1-2 years (Probert and Longley, 1989; McGilp *et al*., 2020). As such, NWR seed cannot be stored in seed banks or repositories unless maintained on an annual basis. NWR is also a monoecious outcrosser with severe inbreeding depression, increasing the difficulty of genetic mapping studies, and requiring the maintenance of effective population sizes for the species survival in natural settings. Currently, genomic resources in the species are limited to studies using a small number of molecular marker studies including isozymes (Lu *et al*., 2005), restriction fragment length polymorphisms (RFLP) (Kennard *et al*., 1999; Kennard *et al*., 2002), simple sequence repeats (SSR) (Kahler *et al*., 2014), and single nucleotide polymorphisms (SNP) (Shao *et al*., 2020). Alignment of molecular markers to a reference genome can more readily provide researchers the ability to investigate the functional relationships between genes and traits of interest, important physiological mechanisms, and the architecture of genetic diversity within the species.

As a recently cultivated crop, the identification and fixation of important domestication traits, such as non-shattering seed phenotypes, is a primary focus of NWR variety development. Although advantageous in natural environments, seed shattering causes significant yield loss in cultivated settings and has been strongly selected against during crop domestication (Doebley, 2006; Fuller *et al*., 2009). In NWR, loss due to shattering can range from 10-20% in a 24-hour period and can be as severe as 70% over a harvest season, the most damaging of which, is the loss of mature seed (Imle, 2001). In cereals, the formation of an abscission layer in the pedicle or rachis is necessary for shattering. While mechanisms to reduce the abscission layer have evolved in different species at different times, the convergent evolution of the non-shattering trait is often the result of independent mutations at orthologous loci in response to strong artificial selection (Doebley, 2006; Purugganan and Fuller, 2009; Lenser and Theißen, 2013; Olsen and Wendel, 2013; Tranbarger *et al*., 2017). This convergence of shattering resistance mechanisms within the grass family has afforded researchers the ability to utilize comparative genomic approaches to identify candidate genes within new species of interest (Van Deynze *et al*., 1998; Nalam *et al*., 2006; Kahler *et al.,* 2014; Fu *et al*., 2019). Initial genetic studies suggest the genetic control of non-shattering in NWR is recessive, putatively controlled by two to three genes, and likely orthologous with several *O. sativa* shattering-related genes (Elliott and Perlinger, 1977; Kennard *et al*., 2000; Kennard *et al*., 2002).

Comparative genomics across the grass species, particularly the cereals, has led to an expansion of knowledge in regard to species’ genome evolution and function. Historically, *O. sativa* has served as a model species for comparative mapping within the grass family given its relatively small genome size and conservation of gene content and relative gene order among the grasses (Zhang *et al*., 2004). As a part of the Oryzeae tribe, members of the *Zizania* genus are considered crop wild relatives of *O. sativa* (Porter, 2019), and techniques including hybridization (Liu *et al*., 1999; Shan *et al*., 2005; Yang *et al*., 2012), protoplast fusion (Liu *et al*., 1999), and gene introduction (Abedinia *et al*., 2000), have been utilized to introgress favorable traits from these species into *O. sativa*. Early comparative mapping studies in NWR revealed significant collinearity with *O. sativa* (Kennard *et al*., 2000; Kahler *et al*., 2014) as well as duplications in the copy number of two *O. sativa Adh* genes (Hass *et al*., 2003). Duplication events have been hypothesized in NWR given the species has three additional chromosomes, in comparison to *O. sativa*, which appear to be duplicates of *O. sativa* chromosomes 1, 4, and 9 (Kennard *et al*., 2000). Comparative analysis between a cultivated *Zizania latifolia* variety and *O. sativa* has also revealed significant collinearity and evidence of a duplication event in *Z. latifolia* ∼10.8-16.1 million years after the two species diverged from one another (Guo *et al.,* 2015).

In this study, a cultivated variety of *Z. palustris,* ‘Itasca-C12’, was chosen for sequencing as it is the most widely grown NWR cultivar in MN and the industry standard for NWR research. From 2016-2017, plants were self-pollinated twice in a greenhouse at the UMN North Central Research and Outreach Center in Grand Rapids, MN to reduce the high level of heterozygosity. Here, we present a chromosome-scale assembly of the NWR genome based on PacBio sequencing as well as Chicago and Hi-C libraries and *ab initio* and evidence-based structural annotation generated using RNA-seq from eight tissues, which will serve as a foundational resource for building a new, modern genomic toolkit for this species. Additionally, we demonstrate the utility of this important resource for both conservation management of natural stands and breeding applications for commercial cultivation.

## MATERIALS AND METHODS

### Plant Materials

In 2018, leaf tissue from Itasca-C12 was collected from a single S2 plant for sequencing. Self-pollinated seed from the individual plant was harvested and stored in water at 3°C in the dark (Oelke and Albrecht, 1978; Oelke and Porter, 2016; McGilp *et al*., 2020). Given that NWR seed is recalcitrant and *ex-situ* seed storage is not currently feasible for this species, seed has not been deposited to a seed bank and is maintained in the UMN NWR breeding, genetics, and conservation program. To preserve the allelic diversity present within the sequenced line, seed is planted annually, and crosses are made between individual plants for use in future studies.

For RNA-seq, 10 Itasca-C12 S3 plants were grown during the spring of 2019 in the UMN Plant Growth Facilities in St. Paul, MN. Eight tissue types (male florets, female florets, leaf, leaf sheath, root, seed, stem, and a whole un-emerged panicle) (Figure S1) were harvested from three individual plants and pooled for sequencing. Leaf, leaf sheath, root, stem, and whole un-emerged panicle tissues were collected at the early boot stage or principal phenological stage (PPS) 41 (Duquette *et al*., 2019). Male and female floret tissues were collected at the end of panicle emergence or PPS 59, and seed was collected when 90% of seed on a panicle was fully ripe or at PPS 89.

### Whole Genome Sequencing and de novo Assembly

Single-plant gDNA (25 µg) was extracted using previously described methods (Zhang *et al*., 1995) and quantified using a Qubit 2.0 Fluorometer (Life Technologies, Carlsbad, CA, USA). Library preparation, sequencing, and assembly were conducted by Dovetail Genomics (Santa Cruz, CA, USA). Sequencing was performed on the Pacific Biosciences (PacBio) Sequel System with eight Single Molecular Real-Time (SMRT) 1M cells to generate 22.6 Gb of sequence data (Table S1). Chicago and Hi-C libraries were prepared as described in Putnam *et al*. (2016) and Lieberman-Aiden *et al*. (2009), respectively. Sequencing libraries were generated using NEBNext Ultra enzymes and Illumina-compatible adapters, and each library was sequenced on an Illumina HiSeqX Ten series platform.

**Table 1.** Summary statistics for PacBio and Dovetail HiRise Assembly with Chicago and Dovetail Hi-C libraries for *Zizania palustris* cultivar, Itasca-C12.

| Metric | Chicago + Dovetail<br>HiRise Assembly | Chicago + Hi-C + Dovetail<br>HiRise Assembly |
| --- | --- | --- |
| Total Length | 1,288.50 Mb | 1,288.77 Mb |
| N50 | 0.689 Mb | 98.770 Mb |
| N90 | 0.170 Mb | 39.126 Mb |
| L50 | 516 scaffolds | 6 scaffolds |
| L90 | 1,928 scaffolds | 14 scaffolds |
| Longest scaffold | 4.8 Mb | 118 Mb |
| Number of scaffolds | 4,834 | 2,183 |
| Number of scaffolds > 1kb | 4,747 | 2,096 |
| Contig N50 | 377.72 kb | 377.56 kb |
| Number of gaps (% of genome) | 3,240 (0.25) | 5,904 (0.27) |

Genome assemblies were performed using the FALCON 1.8.8 pipeline (www.pacb.com) using a length cut-off that corresponded to 50× coverage of data during the initial error-correcting stage. Error-corrected reads were then processed by the overlap portion of the FALCON pipeline. The assembly was polished through PacBio’s Arrow algorithm from SMRT Link 5.0.1 using the original raw reads. Finally, the input assembly, PacBio reads, Chicago library reads, and Hi-C library reads were used as input data for HiRise, a software pipeline designed specifically for using proximity ligation data to scaffold genome assemblies (Putnam *et al*., 2016). An iterative analysis was conducted. First, shotgun and Chicago library sequences were aligned to the draft input assembly using a modified SNAP read mapper (http://snap.cs.berkeley.edu). The separations of Chicago read pairs mapped within draft scaffolds were analyzed by HiRise to produce a likelihood model for genomic distance between read pairs, and the model was used to identify and break putative misjoins, to score prospective joins, and make joins above a threshold. After aligning and scaffolding Chicago data, Dovetail HiC library sequences were aligned and scaffolded following the same method. Shotgun sequences were then used to close gaps between contigs using the PBJelly pipeline with default parameters (English *et al*., 2012).

### Transcriptome Sequencing and Assembly

For each of the eight tissues described above, RNA was extracted using a Qiagen RNeasy kit (product # 74104) and quantified using RiboGreen® RNA quantitation (www.thermofisher.com). RNA-seq library preparations were conducted with a Ribo-Zero® ribosomal RNA (rRNA) reduction. Sequencing (150 bp paired-end reads) was performed by the UMN Genomics Center (UMGC; http://genomics.umn.edu/) on an Illumina NovaSeq S Prime (SP) flow cell (Table S2). Quality scores and potential adapter contaminants were screened using FastQC version 0.11.8 (Andrews, 2010). Low-quality bases and adapter contamination were trimmed using Trimmomatic version 0.33 (Bolger *et al*., 2014). Reads were screened for presence of standard Illumina sequencing adapters, then trimmed based on base quality in sliding windows of 4bp.

**Table 2.** Summary statistics and name designations for the largest 17 scaffolds of the *Zizania palustris* Itasca-C12 genome including chromosome name, original scaffold name, size of the scaffold, and the number of gaps, genes, and SNPs per scaffold at each downsampling step using data from Shao *et al*. (2020).

| Chromosome | Scaffold | Size (Mb) | # of gaps | # of genes | # of SNPs (7M) | # of SNPs (3.5M) | # of SNPs (1.75M) | # of SNPs (0.875M) |
| --- | --- | --- | --- | --- | --- | --- | --- | --- |
| Chr 01 | 13 | 95.4 | 566 | 4,863 | 4,750 | 1,187 | 60 | 0 |
| Chr 02 | 93 | 103.4 | 323 | 4,815 | 4,730 | 1,226 | 18 | 3 |
| Chr 03 | 3 | 58.8 | 274 | 2,859 | 2,651 | 633 | 7 | 3 |
| Chr 04 | 18 | 98.7 | 775 | 2,986 | 2,102 | 558 | 26 | 0 |
| Chr 05 | 1065 | 66.6 | 493 | 2,587 | 2,110 | 488 | 52 | 13 |
| Chr 06 | 48 | 118 | 381 | 7,736 | 9,001 | 2,451 | 119 | 7 |
| Chr 07 | 1063 | 42.6 | 219 | 4,334 | 4,547 | 1,079 | 71 | 1 |
| Chr 08 | 1062 | 75.7 | 263 | 3,539 | 3,409 | 911 | 65 | 8 |
| Chr 09 | 1 | 95.1 | 325 | 2,964 | 2,155 | 537 | 34 | 5 |
| Chr 10 | 70 | 111.4 | 404 | 4,994 | 3,626 | 871 | 57 | 9 |
| Chr 11 | 9 | 63.2 | 234 | 2,539 | 1,766 | 416 | 45 | 0 |
| Chr 12 | 415 | 105.9 | 691 | 4,310 | 3,660 | 999 | 80 | 24 |
| Chr 13 | 1064 | 111.3 | 438 | 5,262 | 4,165 | 1,031 | 54 | 10 |
| Chr 14 | 693 | 24 | 80 | 1,450 | 1,522 | 472 | 24 | 2 |
| Chr 15 | 7 | 39.1 | 150 | 446 | 164 | 43 | 0 | 0 |
| Scf 16 | 51 | 13.8 | 104 | 138 | 136 | 16 | 2 | 1 |
| Scf 458 | 453 | 4.3 | 21 | 7 | 310 | 75 | 4 | 0 |

Reads were trimmed from the 3′ end of the reads until the mean base quality score in a window was at least 15. A high level of rRNA contamination was observed from the FastQC results. A nonredundant collection of rRNA sequences was derived from the SILVA database (Quast *et al*., 2013) using the “dedupe2” tool from the BBTools suite (Bushnell *et al*., 2017). rRNA derived reads were filtered with BBDuk version 38.39 (Bushnell *et al*., 2017). A non-redundant database of ribosomal RNA sequences from the SILVA database (Quast *et al*., 2013) was used to screen for rRNA contamination based on K-mer matching with a K-mer size of 25bp and a maximum edit distance of 1. RNA-seq reads across the eight tissues that passed filtering were used to assemble a single transcriptome with Trinity version 2.8.6 (Grabherr *et al*., 2011; Haas *et al*., 2013) using the “*in silico* read normalization” routine with target coverage set to 200 bp, minimum contig size to 250 bp, and K-mer size to 25 bp.

### Gene Annotation

An interspersed repeat database was created *de novo* from the NWR genome using RepeatModeler 1.0.1. The NWR genome was soft- and hard-masked using the combined RepeatModeler-predicted models and existing RepeatMasker models with RepeatMasker (4.0.5). The abundance of each repeat type was quantified in the R statistical environment version 3.6.0 (R Core Team, 2013). The repeat-masked NWR genome was annotated using the Funannotate 1.5.1 pipeline (Palmer and Stajich, 2018), which uses Augustus (3.2.3) for *ab initio* eukaryotic gene prediction as well as PASA (2.3.3) to refine and correct gene models using RNA-seq evidence (Stanke and Morgenstern, 2005; Haas *et al*. 2003). The Augustus Hidden Markov Models were trained on *O. sativa* Japonica version 1.0.46 (Ensembl release 47) gene features. Genome-aligned RNA-seq reads and Trinity *de novo* assembled full and partial transcripts were provided as RNA-seq read evidence to support the gene prediction process. RNA-seq reads from all NWR tissues were combined and aligned to the NWR genome using STAR 2.7.1 (Dobin *et al*., 2013) using default settings. Completeness of the genic portion of the genome was assessed using the BUSCO version 4.0.0 pipeline (Simão *et al*., 2015). Gene densities as well as the repeat sequences described below were plotted across chromosomes using karyoploteR in R (Gel and Serra, 2017). Blast2GO (Conesa and Götz, 2008) was used to generate functional annotations for the longest protein isoforms based on a BLAST search against the NCBI nr database.

### Genome Evolution and Comparative Genomics

Orthologous gene groups between NWR and 19 other grasses species were identified with OrthoFinder version 2.3.11 (Emms and Kelly, 2019; Table S3). For sequence similarity searches and the generation of trees showing orthologous gene groups, the “BLAST” and “MSA” settings were run in OrthoFinder, respectively. Gene collinearity and syntenic depth between NWR, *O. sativa*, and *Z. latifolia* was evaluated with MCscan using default parameters (Wang *et al*., 2012). The species tree was built with Dendroscope v3 (Huson and Scornavacca, 2012) using the tree file from OrthoFinder as input. We estimated the divergence time between NWR, *Z. latifolia*, and *O. sativa* using the mcmctree program in the PAML (Phylogenetic Analysis by Maximum Likelihood) software package version 4 (Yang, 2007) using a divergence time of 15 million years for *O. glaberrima* and *O. barthii* from *O. sativa* and a tree root age of 30 million years as priors, similar to methods in Guo *et al*. (2015). The whole-genome duplication event was dated by finding the synonymous substitution rate (*KS*) in PAML and converting to geological age using the equation: time (in millions of years) = *KS*/(2×*r*) where *r* is the average *KS* per year (6.5 × 10^-9^ in cereals; Guo *et al*. 2015; Blanc and Wolfe, 2004).

**Table 3.**
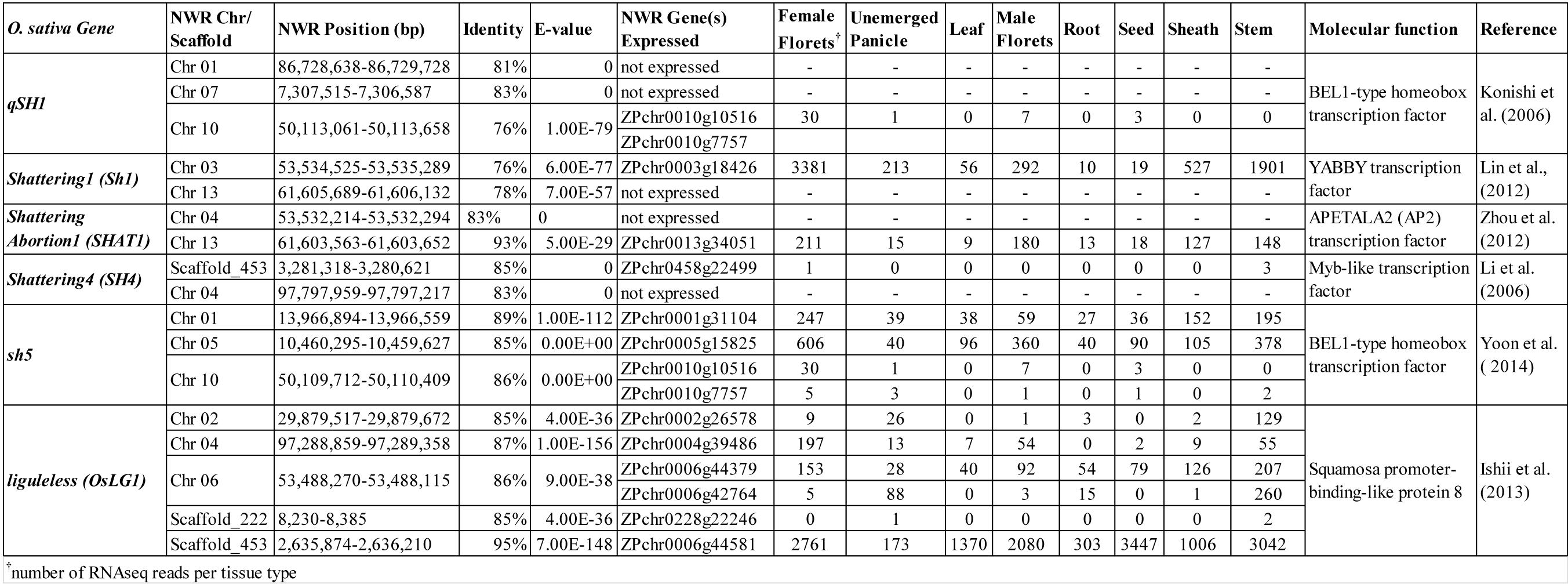
List of *Zizania palustris* orthologs of *Oryza sativa* genes associated with seed shattering and their relative RNA expression levels in *Z. palustris*.

### NWR SNP Distribution and Genotyping-by-Sequencing Read Depth

In order to demonstrate the utility of the NWR reference genome for genetic studies, previously published genotyping by sequencing (GBS) data used to call SNPs without the use of the reference genome (Shao *et al*., 2020) were reanalyzed. The raw sequence data initially reported by Shao *et al*. (2020) can be found at the National Center for Biotechnology Information Short Read Archive (NCBI SRA) under accession number PRJNA574141. FASTQ files were aligned to the genome using the Burrows-Wheeler Aligner (BWA-MEM) version 0.7.17 (Li, 2013) using default parameters. SNPs were called with SAMtools version 1.9 mpileup and BCFtools version 1.2 (Li, 2011) using default parameters through GNU parallel (Tange, 2018). The effect of sequencing depth was also evaluated by sub-sampling the FASTQ files by factors of 2-, 4-, and 8-fold using an awk script to simulate sequencing at lower read depths.

### Identification of NWR Genes Putatively Associated with Seed Shattering

The command line version of BLAST was used to search the NWR genome for orthologs known to be involved with seed shattering resistance in *O*. *sativa* (Konishi *et al*., 2006; Li *et al.,*2006; Lin *et al.,* 2012; Zhou *et al.,* 2012; Ishii *et al.,*2013; Yoon *et al.,* 2014). Candidate selection for NWR shattering genes was improved by comparing our genome annotation with genes submitted to the UniProt database. Genes with measurable expression levels were subsequently checked in different NWR tissues for further validation.

### Code and Data Availability

All *Z*. *palustris* sequencing data generated from this project have been deposited at the NCBI Sequence Read Archive under BioProject PRJNA600525 (Table S1). The whole genome shotgun project has been deposited at the NCBI GenBank under the accession JAAALK000000000. The version described in this paper is JAAALK010000000. Other supporting data have been deposited at the Data Repository for the University of Minnesota (DRUM) under the DOI (10.13020/ha32-4735). All code for the analysis described in this manuscript can be found at https://github.com/UMNKimballLab/NWRGenomeAssembly_v1.0.

## RESULTS AND DISCUSSION

### The Genome Assembly of Northern Wild Rice

In this study, we present the first NWR (*Zizania palustris*) whole genome assembly, which was built using PacBio long-read sequencing and anchored with HiC and Chicago library reads via the HiRise assembly software (www.dovetailgenomics.com). PacBio sequencing of the NWR cultivar, Itasca-C12, generated 7,023,180 reads with a read N50 size of 34 kb (Figure S2). The initial *de novo* assembly using the PacBio FALCON Assembler with default parameters produced 3,689 scaffolds, with an average size of 85 kb and a N50 contig size of 386.6 kb. Chicago library sequencing produced 411 million 2×150 bp paired end reads and provided ∼115× physical coverage of the genome (1-100 kb pairs). HiC library sequencing produced 432 million 2×150 bp paired end reads and provided ∼1,000× physical coverage of the genome (10-10,000 kb pairs). The final HiRise assembly, with HiC and Chicago libraries, consisted of 2,183 scaffolds (L50 = 6 scaffolds; N50 = 98.9 Mb), totaling 1.29 Gb (Table 1; Figure S3). The PBJelly pipeline using default parameters filled only a small fraction of gaps (63 out of 5,904), which totaled ∼0.3% of the genome (Table S4). In comparison, the recent *Z. latifolia* assembly (Guo *et al*., 2015) had a L50 of 305 scaffolds, a N50 of 604.9 kb, and had a total size of 604.1 Mb. One of the many utilities of a sequenced genome is to explore the evolutionary relationships between species. The *Z. latifolia* assembly was the first annotated genome assembly of any *Zizania* species, providing a genome-wide view of these types of relationships for the first time. The inclusion of the NWR genome into these comparisons will help strengthen our understanding of the evolutionary relationships and timeline of the Oryzeae tribe. For example, our assembly demonstrates the size of the NWR genome assembly is ∼400 Mb larger than initial estimates of 860 Mb (Kennard *et al*., 2000), which is ∼3× the size of *O. sativa* (Sasaki, 2005) and ∼2× the size of *Z. latifolia* (586-594 Mb; Guo *et al*., 2015).

To designate chromosome numbers for NWR, we utilized a comparative linkage map of NWR and *O. sati*va (Kennard *et al.,* 2000). Overlaps between the maps and the largest scaffolds were identified, totaling 1.21 Gb in length, across the 15 chromosomes (Table 2). Two additional scaffolds, scaffolds 16 (13.8 Mb) and 458 (4.3 Mb), were quite large and likely represent large unassembled chromosomal fragments or the short arm(s) of a chromosome. Heterozygosity within the sequenced S2 Itasca-C12 plant could have caused high densities of single nucleotide variants and structural variations, such as repeat sequences and coverage gaps, throughout the genome, which may have contributed to the difficulty of integrating scaffolds 16 and 458 with others. Often in such assemblies of heterozygous individuals, homozygous regions of homeologous chromosomes can be collapsed into a single contig, while those of heterozygous regions result in two alternative contigs (Pryszcz and Gabaldón, 2016). Genome assembly software is often unable to resolve those heterozygous alternative contigs, resulting in contigs that cannot be linked and fragmentation of the genomic region. Scaffold 458 is a good example of this phenomenon, as it appears to be a highly heterozygous region of the genome, consisting primarily of coding regions with limited repetitive elements. Despite the two unplaced scaffolds, the NWR genome assembly appears to be largely complete.

Using the resources available to us, namely the close phylogenetic relationship with *O. sativa*, we used comparative analyses to evaluate potential placements for scaffolds 16 and 458. Utilizing the reference genome of a closely -related species during a *de novo* assembly is often used to resolve questions regarding fragmented or misassembled contigs and scaffolds, to orient them along chromosomes, and to provide useful information for genome annotation (Vezzi *et al.,* 2011; Bae *et al.,* 2014; Lischer and Shimizu, 2017). We observed significant collinearity between NWR scaffold 16 and *O. sativa* chromosome 7, which is also collinear with NWR chromosomes 7 and 14 (Figure 1D). Both of these chromosomes have arms ending with dense genic regions (Figure S4). If the genome assembler could not, in fact, combine these scaffolds due to high heterozygosity, this would explain the lack of assembly. For scaffold 458, we observed significant collinearity with *O. sativa* chromosome 4, which is also collinear with NWR chromosomes 4 and 15 (Figure 1D). While scaffold 458 is highly genic, NWR chromosome 4 has dense genic regions at the ends of each arm and chromosome 15 does not have any dense genic regions (Figure S4). We hypothesize scaffold 458 is most likely a part of chromosome 15. While we were able to utilize the Kennard *et al*. (2002) linkage map to help designate NWR chromosomes, the resolution of the map was unable to help resolve these issues and a dense molecular linkage map will be needed in the future.

**Figure 1.**
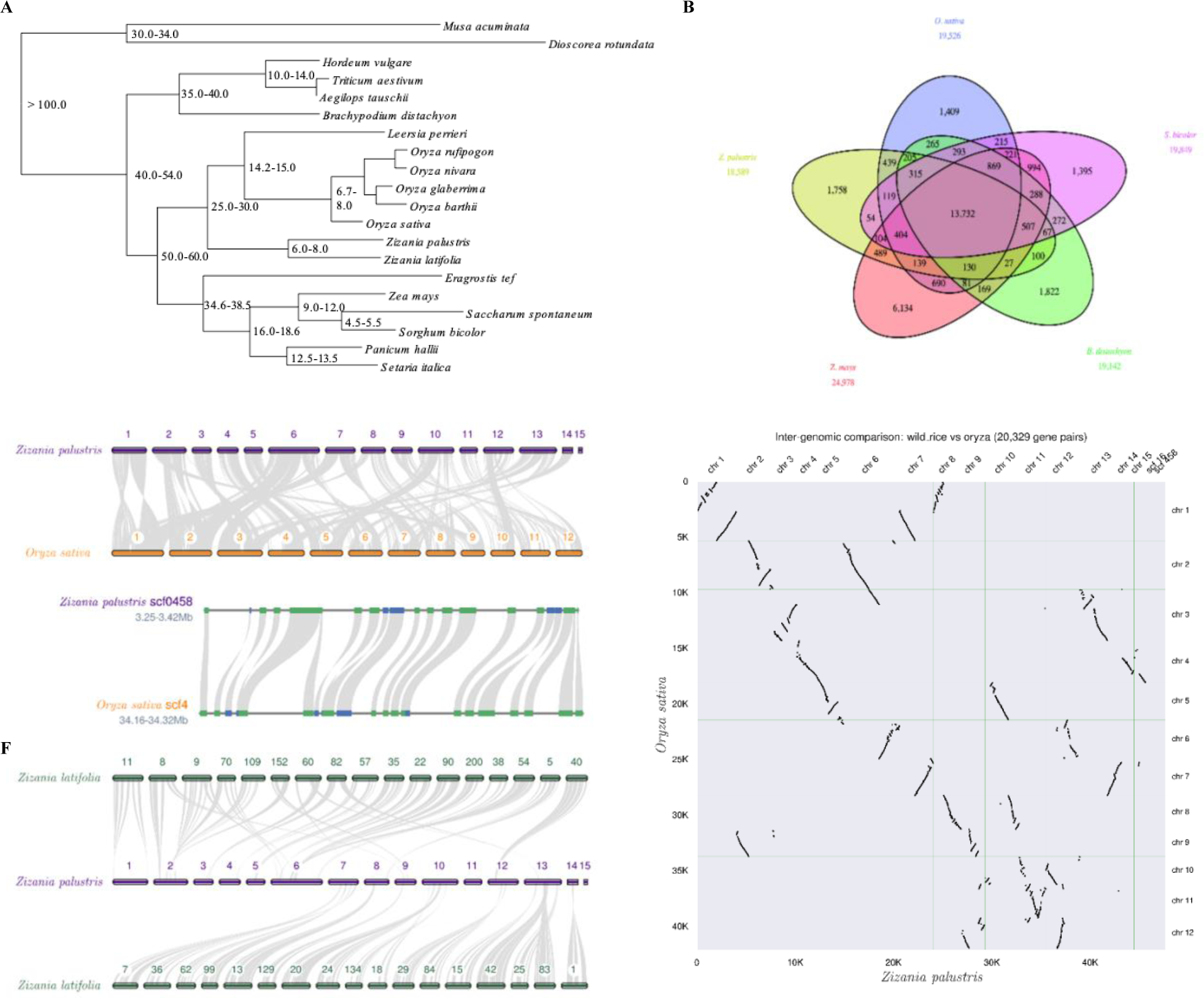
Genome evolution of *Zizania palustris* including: A. A phylogenetic tree of *Zizania palustris* and other *Poaceae* family members using single-copy orthologs. Numbers at nodes represent divergence times in millions of years ago (MYA), B. A venn diagram showing the number of orthogroups for *Oryza sativa*, *Zea mays*, *Sorghum bicolor, Brachypodium distachyon* and *Z. palustris*, C. Synteny between *Z. palustris* and *O. sativa*, D. dot plot showing collinearity between *Z. palustris* and *O. sativa*, E. Microcollinearity between *Z. palustris* and *O. sativa* showing 10 genes on either side of the *sh4* locus and its putative ortholog in *Z. palustris* (green indicates the + strand and blue indicates the - strand), and F. Collinearity between *Z. palustris* and *Z. latifolia*. Panels C-F were all created using MCscan.

While the *O. sativa* genome was able to provide insights into our assembly, we wanted to assess the utility of using the NWR assembly as a guide to help improve the assembly of *Z. latifolia*. The *Z. latifolia* reference genome is largely fragmented, consisting of 761 super-scaffolds (Guo *et al*., 2015). Due to the large number of *Z. latifolia* scaffolds, we compared our NWR assembly only to the largest 34 *Z. latifolia* scaffolds when evaluating collinearity between the two species (Figure 1F). These analyses provided insight into potential alignments of multiple *Z latifolia* scaffolds within a single NWR chromosome. For example, we verified that *Z. latifolia* scaffolds 22, 90, 200, 38, and 54 are syntenic with NWR chromosome 6. Some comparisons, however, demonstrated possible chromosomal rearrangements between the species. For example, *Z. latifolia* scaffolds 8, 9, 11, 70, 152, 60, and 82 all appear to be split between two individual NWR chromosomes. *Z. latifolia* is a diploid with 2 more sets of homeologous chromosomes (*2n*=*2x*=34) than NWR. While we did not evaluate all super-scaffolds for collinearity, it is possible for researchers to now do so with both *Z. latifolia* and *O. sativa* genomes. Hopefully in the near future, the *Z. latifolia* genome assembly can be improved to dissect the relationships between NWR chromosomes and the 2 additional sets in *Z. latifolia*.

### Transcriptome Assembly and Annotation

We utilized eight different tissue types while building the transcriptome assembly. RNA-seq generated 446,755,584 reads across all tissue types, with an average of 55.8 million reads per tissue. The rRNA reduction step prior to sequencing had efficiency issues due to a larger than expected rRNA content, ranging from 6.7-86.4% among tissues (Table S1). Leaf sheath, whole un-emerged panicle, and root tissues had the largest rRNA contamination issues. The Trinity transcriptome assembly, which used filtered reads across all tissues, generated 689,344 transcripts, with an average contig length of 783 bp and a N50 contig length of 1,484 bp (Table S5). There were high levels of heterozygosity within sequenced individuals, as evidenced by the large number of transcripts and total length of the assembly, implying that the separation of alleles and the CD-HIT-EST did not collapse very many transcripts (98% similarity). BUSCO assessment of the transcriptome assembly using 4,896 single-copy Poales orthologues showed that 87.7% of the conserved Poales orthologues were assembled (BUSCO results string C:87.7%[S:25.1%,D:62.6%],F:4.2%,M:8.1%,n:4896). Most of the Poales orthologues that were detected were duplicated, suggesting large numbers of either alternative splicing variants or splitting of allelic variants into separate contigs in this transcriptome assembly.

The annotated genome resulted in 47,696 predicted gene models, of which 46,491 (97.5 %) were putative protein-coding genes. Our annotation was similar to those of *Z. latifolia* and *O. sativa*, which contain 43,703 (Guo *et al.,*2015) and ∼40,000-50,000 (Goff *et al*., 2002; Yu *et al*., 2002) putative protein-coding genes, respectively. The average NWR gene size was 2,905 bp, with an average of 4.6 exons and 3.6 introns. In *Z. latifolia*, the average gene size is 990 bp with a mean of 4.7 exons per gene (Guo *et al*., 2015) and in *O. sativa*, the average gene size is 2,853 bp with a mean of 4.9 exons per gene (Yu *et al*., 2005). Of the 46,491 putative protein-coding genes, gene ontology (GO) terms could be assigned to 24,484 protein-coding genes (52.6%). The most abundant GO terms are depicted graphically in Figure S5 along with their description and abundance in Table S6.

To evaluate the structural and functional features of the NWR genome, coding regions and repetitive elements were characterized. In our whole genome assembly, repetitive elements comprise ∼76.4% of the NWR genome (Figure 2; Figure S4; Table S7)). *Gypsy* and *Copia* retrotransposon superfamilies were the most prevalent (59.2%). The remaining repetitive elements were primarily unclassified elements (10.7% of the genome) and DNA elements (∼5.7%). Long- and short-interspersed retrotransposable elements (LINEs and SINEs) covered ∼0.75% of the genome. The highly repetitive nature of the NWR genome is consistent with the majority of plant genomes, where the expansion, loss, and movement of these elements have played key roles in genome and chromosome evolution (Uozu *et al*., 1997; Kubis, 1998; Feuillet and Keller, 2002; Mehrotra and Goyal, 2014). Approximately 50% of the *O. sativa* genome is comprised of repetitive sequences (Kurata *et al*., 1994) and the expansion of repetitive elements in NWR appears to be one of the causes of its large genome size, relative to *O. sativa*. A comprehensive structural analysis of the repetitive elements in this NWR reference assembly would provide more valuable insights into the evolution of NWR and its relationships in the *Zizania* genus and *Oryza* tribe.

**Figure 2.**
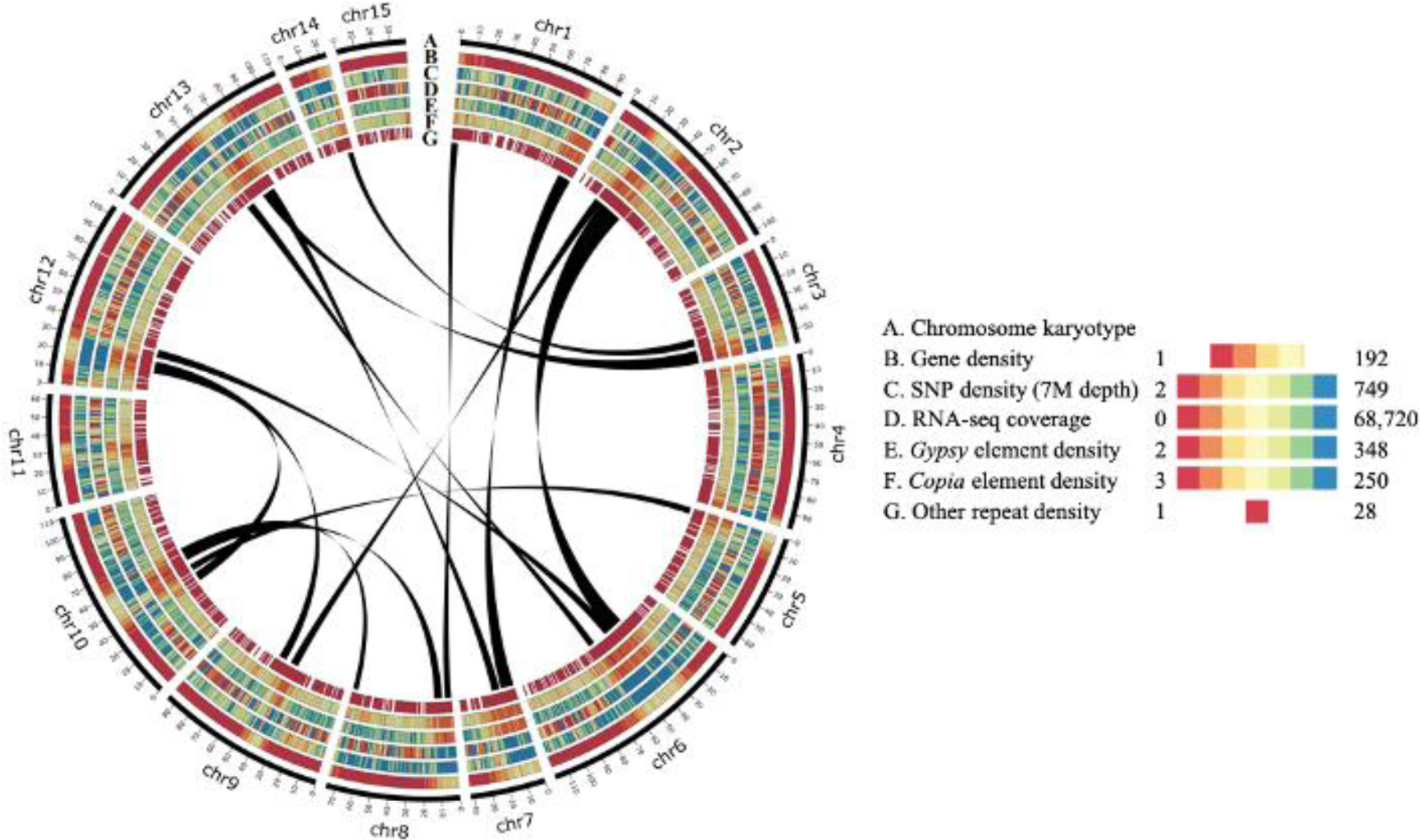
The genome landscape of *Zizania palustris*. Circos-plot circles represent: A. Assembled chromosomes (scale in megabases), B. Gene density, C. SNP density at 7M read depth, D. RNA-seq coverage, E. *Gypsy* element repeat density, F. *Copia* element repeat density, and G. Other repetitive element density. Links between chromosomes depict synteny of gene blocks between chromosomes.

### Inclusion of NWR in Poaceae Orthology Analyses Confirms Phylogenetic Relationships

A cornerstone of comparative genomics is the characterization and comparison of orthologous and paralogous genes across species of interest, providing informative insights into their evolutionary relationships. Orthologous genes, in particular, have been widely characterized across Poaceae, especially among crop species, and have been instrumental in the characterization of the significant collinearity identified within the family (Kellog and Watson, 1993; Bennetzen and Freeling, 1997; Devos and Gale, 1997; Gaut, 2002; Schnable *et al.,* 2012). Despite the rapid increase in our understanding of such familial relationships, numerous species within the family still have extremely limited genomic resources, which impedes our understanding of the evolutionary relationships within the family. Even within Oryzeae, a widely researched tribe with a distinct monophyletic lineage (Kellogg and Watson, 1993), the taxonomic separation of monoecious and bisexual genera, based on morphological and reproductive data, into Oryzinae and Zizaniinae subtribes (Hitchcock and Chase, 1951; Stebbins and Crampton, 1961) was initially disputed for decades (Terrell and Robinson, 1974; Duvall *et al*., 1993; Ge *et al*., 2001). However, the characterization of sequences such as *adh* and *matK* genes (Ge *et al*., 2002; Xu *et al*., 2008; Xu *et al*., 2010) have helped to confirm this taxonomic classification. In this study, we present a phylogenetic analysis, based on the protein-coding orthogroups of 20 species in the grass family that is consistent with previous findings supporting the placement of NWR in the tribe Oryzeae and subtribe Zizaniinae (Figure 1A) (Duvall *et al*., 1993; Ge *et al*., 2002; Tang *et al*., 2010).

A large number of shared and divergent orthogroups were identified between NWR and several major grass species including *O. sativa*, *S. bicolor*, *Z. mays*, and *B. distachyon*. *Z. mays* (Figure 1B). A total of 13,732 orthogroups were shared between all five species, which is consistent with other studies evaluating the distribution of shared gene families in Poaceae (International Brachypodium Initiative, 2010; Carballo *et al.,* 2019). *Z. mays* had the largest number of unique orthogroups (6,134) amongst the five species, possibly due to the large divergence time between Oryzoideae and Panicoideae subfamilies or the large pan-genome size of *Z. mays*, which has a considerable number of dispensable genes (Hirsch *et al.,* 2014). Evaluation of the clustering of orthogroups within the Oryzeae tribe alone revealed 14,120 orthogroups shared between NWR, *Z. latifolia*, *O. sativa*, *O. rufipogon*, and *O. glaberrima* (Figure S6). NWR had the most unique orthogroups (1,731), compared to only 538 and 712 orthogroups classified in *Z. latifolia* and *O. sativa*, respectively. These unique orthogroups may be attributed to specific adaptive characteristics within the species. For example, NWR and *Z. latifolia* have diverged significantly in their growth habits. NWR is an annual plant, adapted to colder climates, while *Z. latifolia* is a perennial, adapted to warmer climates. Cultivated *Z. latifolia* is also unique, as it is persistently colonized with a fungal endophyte, *Ustilago esculenta*, which has resulted in edible stems and the loss of flowering (Yu, 1962; Chans and Thrower, 1980).

The NWR Genome is Highly Collinear with *O. sativa*

Comparative analyses among members of *Oryzeae* is of particular interest to *Zizania* researchers, given their close phylogenetic relationships and the wealth of scientific knowledge available within the tribe. We estimated that NWR diverged from *O. sativa* ∼25 MYA (Figure 1A), which is consistent with previous estimates (Tang *et al.,* 2010; Guo *et al*., 2015). Comparative analysis between the two species’ genomes revealed a picture of collinearity conserved on both the macro and micro levels, along with duplications, chromosomal reshuffling, and inversions, indicative of speciation and whole genome duplication (WGD) events. We first established that there is a high degree of synteny between the genomes of *O. sativa* and NWR (Figure 1C; Figure 1D). For example, NWR chromosomes 1-3 were highly collinear with *O. sativa* chromosomes 1-3, respectively (Figure 1C). Numerous chromosomal arms of *O. sativa* were shuffled and duplicated within the NWR genome. For example, the individual arms of chromosome 5 of *O. sativa* were split between NWR chromosomes 5 and 10. Similarly, the arms of chromosome 9 of *O. sativa* were split between NWR chromosomes 2 and 9. The largest NWR chromosome, chromosome 6, was an amalgamation of large swaths of *O. sativa* chromosomes 2, 3, and 6. Large-scale chromosomal inversions were also identified on nearly every NWR chromosome/scaffold, which were commonly located at the transition between dense LTR and genic regions (Figure 1D; Figures S4A and D). Inversions are common throughout the plant kingdom and have been characterized broadly in crops and wild relatives across the Solanaceae, Poaceae, and Brassicaceae families (Huang and Rieseberg, 2020). Large-scale inversions, like we see in the NWR genome, are frequently characterized as drivers of speciation and adaptive change (Kirkpatrick and Barton, 2006; Feder and Nosil, 2009; Fuller *et al*., 2018) and may have led to reproductive barriers between *O. sativa* and NWR (Figure 1D). Despite all this variation and genome shuffling, micro-collinearity or gene order within these larger syntenic regions was also observed, as exemplified by the genic region surrounding the *Shattering 4* (*SH4*) locus shown in Figure 1E.

Genome-Wide Comparisons with *Z. latifolia* reveal a Rapid Expansion of Repetitive Elements in NWR

With our new assembly in hand, we were able to compare genome-wide characteristics and relationships between two *Zizania* species for the first time. First, we estimated that the species diverged from one another ∼6.0-8.0 MYA, or ∼17-19 million years after the genera *Zizania* diverged from *Oryza* (Figure 1A). The first estimates of divergence time between NWR and *Z. latifolia* were dated to 3.74 MYA, based on the phylogenic analysis of seven genes, including *Adh*1*a* (Xu *et al*., 2010). Further comparisons between NWR and *Z. latifolia* revealed two genomes with comparable protein-coding genes, 46,491 in NWR and 43,703 in *Z. latifolia*. Guo *et al*. (2015) identified that 4.6% or 2,010 of protein-coding genes in the domesticated *Z. latifolia* genome were lost or carried loss-of-function mutations. The majority of these mutations were involved in plant immunity networks and were most likely due to the persistent *Ustilago* infection. In contrast, the repetitive regions constituted 76.4% (924.4 Mb) of the NWR genome assembly and only 37.7% (227.5 Mb) of the *Z. latifolia* assembly (Guo *et al*., 2015). *Gypsy* and *Copia* elements, specifically, make up a significant portion of the repetitive regions in both species’ genome assemblies (Table S7; Figure S4 B-D; Guo *et al*., 2015). These LTR retrotransposons can impact genomes in a number of significant ways, including variation in genome size within angiosperms (Bennetzen, 2002), regulation of gene networks (Struder *et al*., 2011; Yang *et al*., 2012), and structural changes (Bennetzen *et al*., 2005; Vitte and Panaud, 2005). Studies have estimated that LTR expansion in some species has been relatively rapid. Within the last 6 million years, for example, the arrival and amplification of retrotransposons in maize have effectively doubled the species’ genome size (SanMiguel and Bennetzen 1998; SanMiguel *et al.,* 1998). The same is true for select members of the genus *Oryza*, such as *Oryza australiensis*, which has undergone a recent burst of LTR-retrotransposons in the past three million years (Piegu *et al*., 2006). The large increase of LTRs in the NWR genome, seemingly after the NWR-*Z. latifolia* speciation event 6-8 MYA, suggest this is also true for NWR.

### The NWR Genome Assembly Confirms a Whole Genome Duplication in *Zizania*

Whole genome duplications (WGD) are common in the plant kingdom and have been well documented across the grass family (Paterson *et al*., 2004; Yu *et al*., 2005; Salse *et al*., 2008). Guo *et al*. (2015) identified a WGD event in *Z. latifolia* that occurred ∼10.6-15.9 MYA, which is an estimated 10.8-16.1 million years after the *Zizania*-*Oryza* speciation event. Our study identified a considerable amount of evidence to support that this WGD event also occurred in NWR. To start, the NWR genome is ∼3× the size of the *O. sativa* genome and has twice as many 2:1 orthologue groups, indicating a significant amount of gene duplication (Table S8). The mean length of syntenic blocks in NWR was 2× the length of those in *O. sativa* (Table S9). Additionally, the MCscan dot plot provided excellent visualization of the duplication of every *O. sativa* chromosomal arm within the NWR genome (Figure 1D). We also evaluated syntenic depth between the two species or the number of syntenic regions in the target genome for any given query position (Tang *et al*., 2012) to itemize how many genes were covered in 1-, 2-, to *x*-fold regions. This analysis is more accurate than an orthologue ratio analysis for evaluating large-scale genomic events, such as WGDs, because it is not influenced by small-scale changes, such as tandem duplications or expansions/contractions (Tang *et al.,* 2015). We identified a 2:1 synteny pattern between NWR and *O. sativa*, where 56% of NWR regions had a syntenic depth of 2, or two syntenic blocks per *O. sativa* gene (Figure 3A). Only 5% of *O. sativa* syntenic regions had a syntenic depth of 2. There was no such 2:1 ratio observed between NWR and *Z. latifolia* (Figure 3B). A 2:1 syntenic pattern is often a result of co-orthologous regions driven by large-scale events, such as WGDs (Tang *et al*., 2015), which further supports the hypothesis of a WGD in the *Zizania* genus.

**Figure 3.**
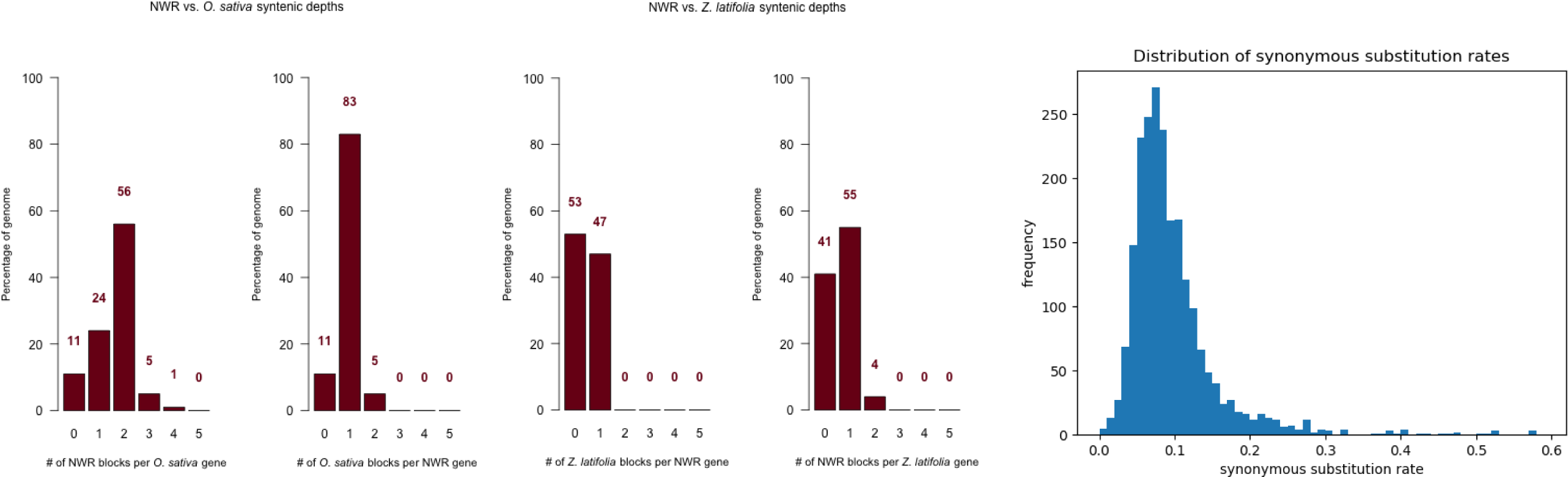
Comparative analyses between northern wild rice (NWR; *Z. palustris*), *O. sativa*, and *Z. latifolia* including A. The distribution of synteny blocks in NWR and *O. sativa* for each *O. sativa* and NWR gene, respectively; B. The distribution of synteny blocks in NWR and *Z. latifolia* for each *Z. latifolia* and NWR gene, respectively; and C. The distribution of synonymous substitution rates (*Ks*) within NWR used to estimate the age of the WGD event in *Zizania*.

The syntenic depth analysis between NWR and *Z. latifolia* was not as informative due to the large number of scaffolds in the *Z. latifolia* genome. In MCscan, we used the default number of 30 or more syntenic genes to establish syntenic blocks between NWR and *O. sativa* (Figure 1C) but had to reduce that number to 10 in order to detect synteny between NWR and *Z. latifolia* (Figure 1F). When the default number of minimum syntenic genes was used, no synteny between NWR and *Z. latifolia* was found. Additionally, the comparisons of the mean block lengths in NWR vs. *O. sativa* and NWR vs. *Z. latifolia* were 6.4 Mb vs.

3.1 Mb and 6.4 Mb vs. 0.2 Mb, respectively (Table S9). The number of gene pairs per block was 135 for NWR vs. *O. sativa* but only ∼15 for NWR vs. *Z. latifolia*. The low number of gene pairs per block between NWR and *Z. latifolia* is most likely a product of the fragmented nature of the *Z. latifolia* genome assembly, rather than a biological observation.

During the calculations of divergence time estimates between NWR and *Z. latifolia*, it initially appeared that the WGD event in NWR followed the NWR-*Z. latifolia* speciation event. Our estimates indicated that the speciation event occurred ∼6.0-8.0 MYA (Figure 1A) and the WGD event ∼0.7-1.7 million years later (∼5.3 MYA) (Figure 3C). This is ∼2.6-9.9 million years later than the *Z. latifolia* WGD event (∼10.6-15.9 MYA), estimated by Guo *et al*. (2015). While this could help explain why the *Z. latifolia* assembly is 589 Mb (Guo *et al*., 2015), or approximately half, the size of the NWR genome (1,290 Mb), we did not identify further evidence to support a second NWR-specific WGD within the *Zizania* genus. This suggests that the resolution of the molecular clock time within this study was not sufficient to resolve the relationship between speciation and WGD events in the *Zizania* genus. Issues using the molecular clock as a technique to infer the dates of major species divergence events have been noted across the plant kingdom as the evolutionary rate of change is often not constant between species or even across a genome (Robinson and Robinson, 2001). These rates can be influenced by a range of factors including life-history traits (Kumar, 2005) and certain evolutionary events, such as rapid radiations or the rapid increase in taxonomic diversity resulting from elevated rates of speciation (Benton, 1999). Fossil records are often used to validate or challenge molecular clock estimates but few *Zizania* fossil records exist (Lee *et al*., 2004; Yost *et al*., 2013) and none have been used to date the NWR speciation event.

Contrary to our initial calculations of divergence, we did identify evidence to support the hypothesis that the WGD event in NWR occurred prior to the NWR-*Z. latifolia* speciation event. The increase in size of the NWR genome in comparison to *Z. latifolia* was associated with an expansion of LTR repetitive elements in NWR, not the coding regions, which were similar in size between the two species (376.5 Mb in *Z. latifolia* vs 304.5 Mb in NWR). Variation in genome sizes in the plant kingdom has long been known to be due to mostly repetitive DNA (Flavell *et al*., 1974). Indeed, genome size doubling due to retrotransposons has been observed in many species, including *O. australiensis* (Piegu *et al*., 2006). Even if species are closely related, they can still differ greatly in their genome sizes after episodes of lineage-specific expansion (Grover and Wendel, 2010). Finally, variation in the number of 2:1 orthologue groups between the species was minimal (Table S8) and the analysis of syntenic depth between them did not reveal a 2:1 pattern (Figure 3B). This evidence collectively supports the WGD event happened prior to the NWR-*Z. latifolia* speciation event.

### Leveraging the Annotated NWR Reference Genome for Plant Improvement

Reliable reference genomes are useful for genetic studies as they can provide insights into evolutionary events and relationships, functional genomic and linkage disequilibrium analyses, and the identification of genes responsible for traits of interest. To highlight the utility of the NWR genome, we re-examined a set of NWR SNPs reported in Shao *et al*. (2020), and identified putative seed-shattering genes in NWR based on the well-characterized shattering genes in *O. sativa* (Konishi *et al.,* 2006; Li *et al*., 2006; Lin *et al.,* 2012, Zhou *et al.,* 2012; Ishii *et al.,* 2013; Yoon *et al*., 2014). We then calculated the number of SNPs within 1Mb of putative shattering genes to assess GBS-derived SNP densities surrounding these genic regions.

In 2020, a small GBS-driven SNP identification study was published to evaluate SNP densities at four GBS read depths for future use in genetic studies (Shao *et al.,* 2020). Here, we present the alignment of the original GBS data (7M reads/sample) to the genome assembly along with sub-sampled sets (∼3.5M, 1.75M, and 0.875M reads) to assess SNP frequency and distribution across the NWR genome (Figure S7). SNP densities decreased drastically when down sampled to less than 3.5M reads with an average distribution of 41.4, 10.6, 0.6, and 0.1 SNPs per Mb at sequencing levels of 7M, 3.5M, 1.75M, and 0.875M reads, respectively (Figure S7). SNPs were also plotted in 1 Mb bins to evaluate their distribution across the genome. With 7M reads, SNP density was highest (up to 400 SNPs/Mb) in gene-rich regions and typically lowest in LTR-rich regions (Figure 2, Figure S7). This pattern was identified across sequencing levels and most chromosomes (Table 2). Gene-poor chromosome 15 and scaffold 16, had the lowest SNP densities, with no bin exceeding 40 SNPs/Mb with 7M reads. Collectively, these results indicate that generation of 3.5M GBS reads or greater is likely necessary for molecular studies in NWR. The restriction enzymes, *Btg1* and *Taq1*, were chosen for Shao *et al* (2020) based on a previous estimate of the NWR genome size (600-800 Mb; Kennard *et al.,* 2000), which was considerably lower than the size of the reference assembly (1.29 Gb). *In silico* digestion of the reference assembly revealed that a *Mst1* and *Pst1* restriction enzyme combination would yield a larger number of SNPs for future GBS studies using the RestrictionDigest perl module (https://metacpan.org/pod/release/JINPENG/RestrictionDigest.V1.1/lib/RestrictionDigest.pm) with default parameters. Plant genetics and genomics are central to plant improvement strategies commonly used by plant breeders to produce new cultivars that are higher yielding, agronomically uniform and pest resistant. The genomics age, in particular, has expanded the possibilities for novel trait discovery in niche crops, like NWR, far beyond those attainable through first-generation molecular markers. For example, we are now able to utilize comparative genomic approaches to identify putative genes associated with important traits of interest in NWR, such as seed shattering, a primary focus in NWR cultivar development. In this study, we queried six *Oryza* shattering-related genes against the NWR genome assembly using BLAST to identify putative genes of interest. Most notably, we identified the ortholog of the *SH4* locus, ZPchr0458g22499 on scaffold 458 (Table 3), a major regulator of abscission layer formation in *O. sativa* (Li *et al*., 2006). This gene was previously identified as a potential seed shattering-related candidate using a NWR linkage map (Kennard *et al*., 2002). Other notable NWR genes include orthologs of *qSH1* (Konishi *et al*., 2006), *Sh5* (Yoon *et al*., 2017), *Shattering1* (Lin *et al*., 2007), *Shattering Abortion1* (Zhou *et al*., 2012), and *OsLG1* (Ishii *et al*., 2013) (Table 3).

Multiple BLAST hits were identified in NWR for eac*h O. sativa* shattering gene we evaluated (twenty hits total for six *O. sativa* genes), indicating that gene duplication may be common across the genome. This is rather likely given the rapid expansion of retrotransposons across the NWR genome and the recent WGD event in *Zizania*, both of which are common causes of gene duplication in plant species (Krasileva, 2019). Examples of duplicated regions harboring putative shattering genes can be visually identified utilizing both the assembly circus plot (Figure 2) and the *O. sativa* collinearity dot plot (Figure 1D). While the expression of several of these paralogous hits was not identified during the analysis of RNA-seq data, which would have further validated gene candidates, it is very possible that the time of tissue collection was not appropriate to capture expression and further testing is needed (Table 3). Several of these candidate NWR shattering loci also co-localized with one another indicating potential clusters of shattering-related genes. For example, we identified the co-localization of two *SH4* candidates on NWR chromosome 4 and two *OsLG1* candidates on scaffold 458. We also identified the co-localization of candidates for *qSH1* and *Sh5*, which are homologous with one another in *O. sativa* (Yoon *et al*., 2017), a phenomenon that is known to happen amongst shattering genes across the grass family (Di Vittori *et al.,* 2019). Comparison of the size of these orthologs revealed that many are of similar size, however a few orthologs in NWR were almost twice the size of *O. sativa* genes (Table S10). Previous studies have identified that differences in gene size can be caused by increases in the amount of intergenic transposable elements (Bennetzen and Ma, 2003; Swigonova *et al*., 2005) as well as duplication events, where one paralog is free from selection resulting in either a loss of function or the development of novel functions within the genome (Panchy *et al*., 2016)

To conclude these initial evaluations, we counted SNPs from Shao et al (2020) within 1 Mb up- and downstream (2 Mb total window size) from the start position of each putative NWR shattering gene (Table 3). Among the 17 largest scaffolds at a read depth of 7M, the number of SNPs ranged from 54 SNPs for the *sh5* candidate ZPchr0001g31104 to 489 SNPs for the *OsLG1* candidate ZPchr0006g44369 (Table S10) with an average number of 254 SNPs surrounding each of the candidate regions. At 3.5M, 1.75M, and 0.875M reads, the average SNP number surrounding the candidate regions was 65, 5, and 1, respectively. It is important to note that these numbers are likely over-estimates of reliable SNPs due to the limited number of samples (8) in the Shao *et al*. (2020) dataset where assessments of minor allele frequencies were negligible. While linkage disequilibrium (LD) has yet to be evaluated in NWR, we suspect LD decays rather rapidly given the species out-crossing habit, which will require a large number of SNPs distributed along the genome to identify causal variants. In maize for example, LD decays at a rate of 1-10kb depending on the chromosome (Yan *et al*., 2009) and large SNPs sets are required in the species. Nevertheless, this examination demonstrates that variation exists to develop assays such as Kompetitive Allele-Specific PCR (KASP) markers to select for favored alleles at these loci (Semagn *et al*., 2013).

## CONCLUSIONS

The NWR genome presented here is an important resource for the advancement of genomic research in this species as well as comparative genomic studies with *O. sativa* and *Z. latifolia*. This *de novo* reference assembly is largely complete, highly repetitive, and 1.5-2x larger than anticipated. The expansion of retrotransposons within the genome and a whole genome duplication prior to the *Zizania-Oryza* speciation event is likely to have led to an increase in the genome size of NWR in comparison with both *O. sativa* and *Z. latifolia*. Both events depict a genome rapidly undergoing change over a short evolutionary time providing new insights into the evolutionary history of the Oryzeae tribe and the grass family in general. The significant collinearity between NWR and *O. sativa* provides NWR researchers with a rich genomic resource to aid in the identification of genes of agronomic importance and provides a unique opportunity to study the genetics of the domestication process in real time.

## Acknowledgements

The authors would like to thank the staff at the University of Minnesota Genomics Center (UMGC) and acknowledge the Minnesota Supercomputing Institute (MSI) at the University of Minnesota for providing resources that contributed to research results reported in this paper. This work was supported by the Minnesota Cultivated Wild Rice Council and by the State of Minnesota, Agricultural Research, Education, Extension, and Technology Transfer program.

---

**Supporting Table S1.** Information for raw PacBio, Illumina, and RNA-seq sequencing data submitted to the National Center for Biotechnology Information Short Read Archive (NCBI SRA) as well as assembly and scaffolding files for both northern wild rice (NWR; *Zizania palustris*) whole genome and transcriptome assemblies. The files can be found under BioProject number PRJNA600525 and BioSample number SAMN13825534.

| File name | Identity | NCBI Accession number |
| --- | --- | --- |
| DTG-DNA-358_cell1.fastq.gz | PacBio SMRT cell 1 | SRR11927429 |
| DTG-DNA-358_cell2.fastq.gz | PacBio SMRT cell 2 | SRR11927429 |
| DTG-DNA-358_cell3.fastq.gz | PacBio SMRT cell 3 | SRR11927429 |
| DTG-DNA-358_cell4.fastq.gz | PacBio SMRT cell 4 | SRR11927429 |
| DTG-DNA-358_cell5.fastq.gz | PacBio SMRT cell 5 | SRR11927429 |
| DTG-DNA-358_cell6.fastq.gz | PacBio SMRT cell 6 | SRR11927429 |
| DTG-DNA-358_cell7.fastq.gz | PacBio SMRT cell 7 | SRR11927429 |
| DTG-DNA-358_cell8.fastq.gz | PacBio SMRT cell 8 | SRR11927429 |
| DTG-HiC-690_R1_001.fastq.gz | Hi-C library | SRR13562678 |
| DTG-HiC-690_R2_001.fastq.gz |  |  |
| DTG-CHI-577_R1_001.fastq.gz | Chicago library | SRR13562677 |
| DTG-CHI-577_R2_001.fastq.gz |  |  |
| Female_S1_R1_001.fastq.gz | Female floret | SRR12661001 |
| Female_S1_R2_001.fastq.gz |  |  |
| Flower_S8_R1_001.fastq.gz | Whole un-emerged panicle | SRR12661000 |
| Flower_S8_R2_001.fastq.gz |  |  |
| Leaf_S2_R1_001.fastq.gz | Leaf | SRR12660999 |
| Leaf_S2_R2_001.fastq.gz |  |  |
| Male_S4_R1_001.fastq.gz | Male floret | SRR12660998 |
| Male_S4_R2_001.fastq.gz |  |  |
| Root_S5_R1_001.fastq.gz | Root | SRR12660997 |
| Root_S5_R2_001.fastq.gz |  |  |
| Seed_S6_R1_001.fastq.gz | Seed | SRR12660996 |
| Seed_S6_R2_001.fastq.gz |  |  |
| Sheath_S3_R1_001.fastq.gz | Sheath | SRR12660995 |
| Sheath_S3_R2_001.fastq.gz |  |  |
| Stem_S7_R1_001.fastq.gz | Stem | SRR12660994 |
| Stem_S7_R2_001.fastq.gz |  |  |

**Supporting Table S2.**
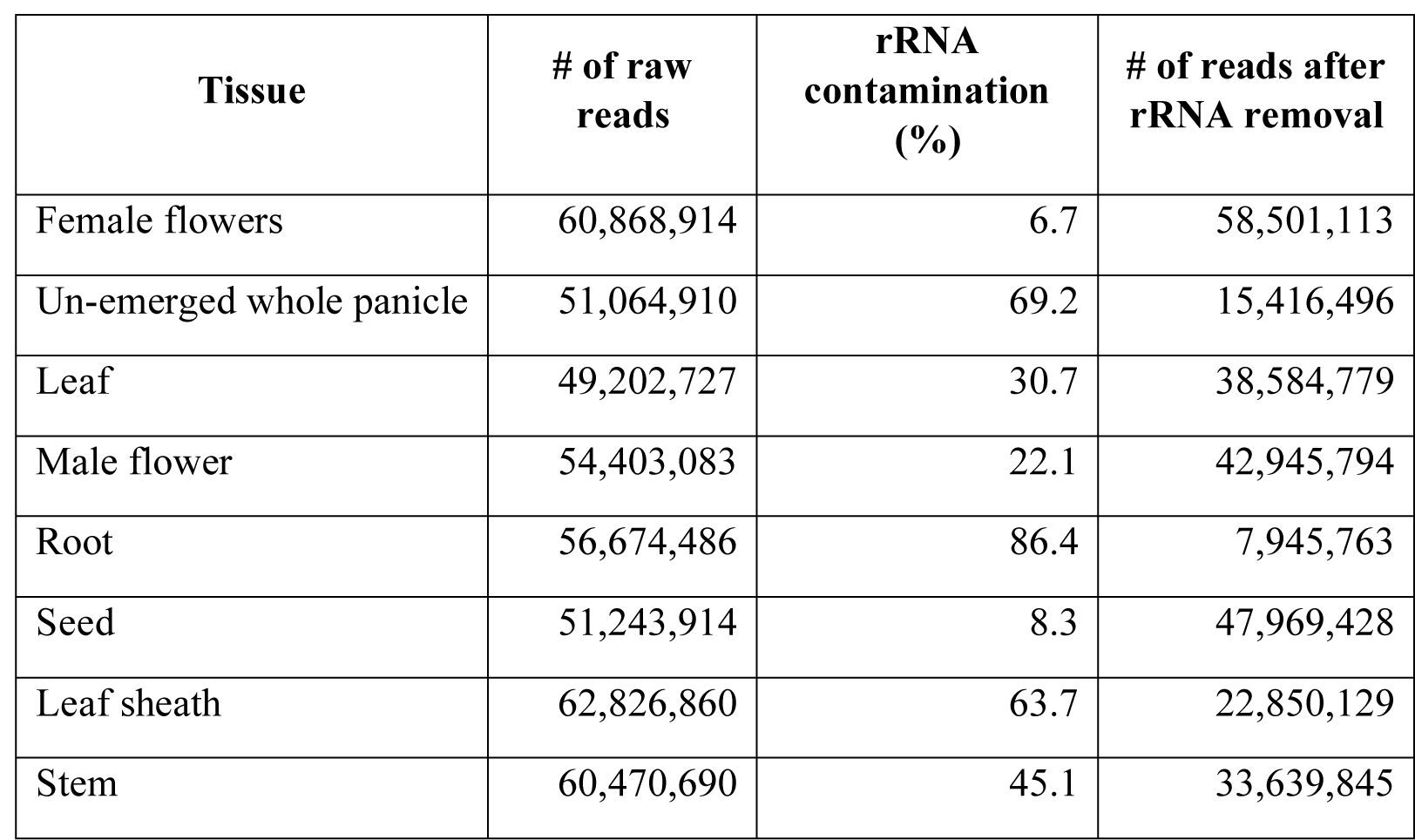
Summary statistics of northern wild rice (NWR; *Zizania palustris*) cultivar Itasca-C12 RNA-seq results including ribosomal RNA (rRNA) contamination.

| **Tissue** | **# of raw reads** | **rRNA contamination (%)** | **# of reads after rRNA removal** |
| --- | --- | --- | --- |
| Female flowers | 60,868,914 | 6.7 | 58,501,113 |
| Un-emerged whole panicle | 51,064,910 | 69.2 | 15,416,496 |
| Leaf | 49,202,727 | 30.7 | 38,584,779 |
| Male flower | 54,403,083 | 22.1 | 42,945,794 |
| Root | 56,674,486 | 86.4 | 7,945,763 |
| Seed | 51,243,914 | 8.3 | 47,969,428 |
| Leaf sheath | 62,826,860 | 63.7 | 22,850,129 |
| Stem | 60,470,690 | 45.1 | 33,639,845 |

**Supporting Table S3.**
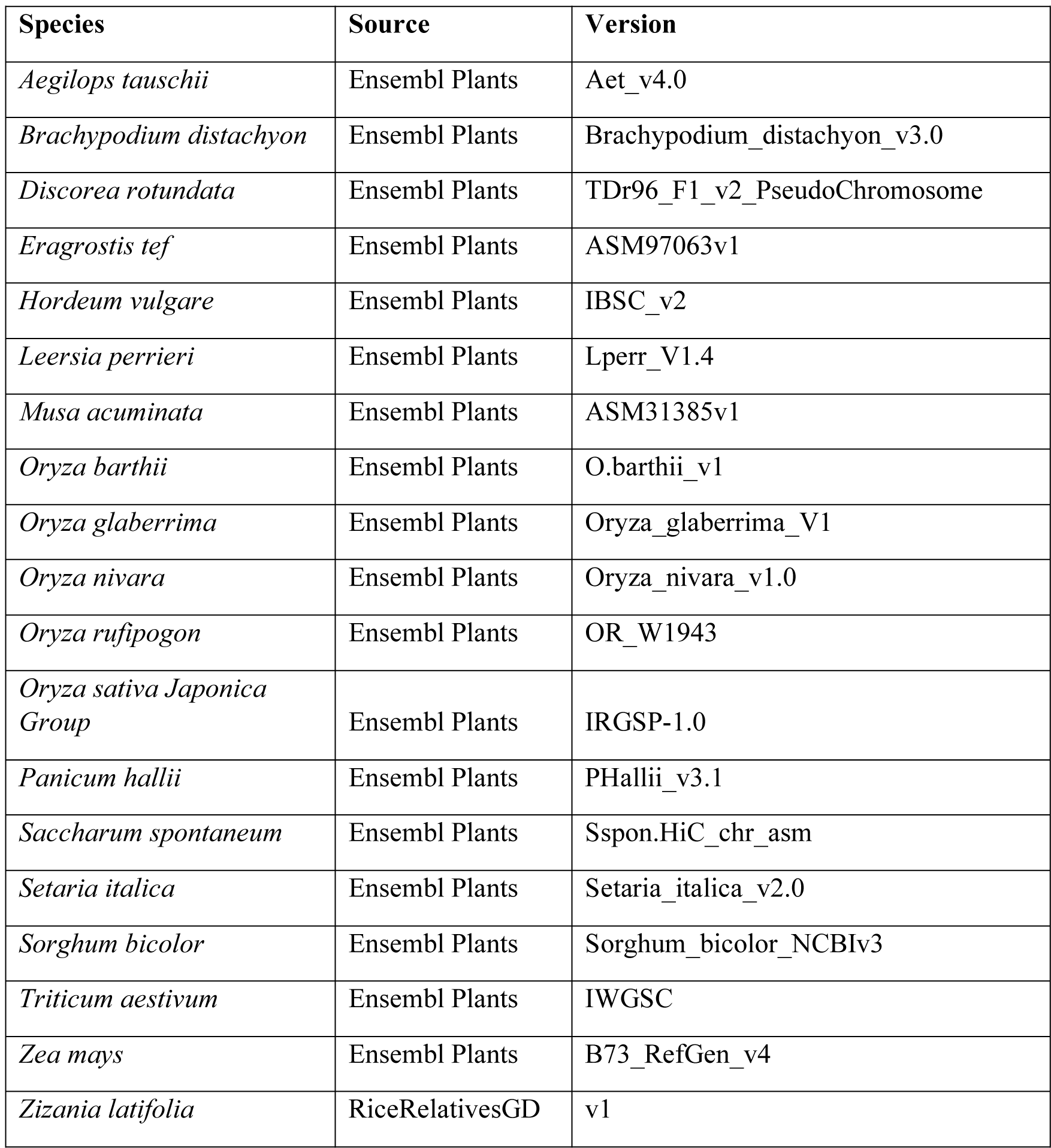
List of grass species included in the OrthoFinder gene group analyses. All 20 species (including NWR) were used in the analysis to generate the species tree in Figure 2A, but independent analyses used to generate the venn diagrams in Figure 2B and Supporting Figure S7 were performed with only the species depicted in each respective figure.

| Species | Source | Version |
| --- | --- | --- |
| *Aegilops tauschii* | Ensembl Plants | Aet_v4.0 |
| *Brachypodium distachyon* | Ensembl Plants | Brachypodium_distachyon_v3.0 |
| *Discorea rotundata* | Ensembl Plants | TDr96_F1_v2_PseudoChromosome |
| *Eragrostis tef* | Ensembl Plants | ASM97063v1 |
| *Hordeum vulgare* | Ensembl Plants | IBSC_v2 |
| *Leersia perrieri* | Ensembl Plants | Lperr_V1.4 |
| *Musa acuminata* | Ensembl Plants | ASM31385v1 |
| *Oryza barthii* | Ensembl Plants | O.barthii_v1 |
| *Oryza glaberrima* | Ensembl Plants | Oryza_glaberrima_V1 |
| *Oryza nivara* | Ensembl Plants | Oryza_nivara_v1.0 |
| *Oryza rufipogon* | Ensembl Plants | OR_W1943 |
| *Oryza sativa Japonica Group* | Ensembl Plants | IRGSP-1.0 |
| *Panicum hallii* | Ensembl Plants | PHallii_v3.1 |
| *Saccharum spontaneum* | Ensembl Plants | Sspon.HiC_chr_asm |
| *Setaria italica* | Ensembl Plants | Setaria_italica_v2.0 |
| *Sorghum bicolor* | Ensembl Plants | Sorghum_bicolor_NCBIv3 |
| *Triticum aestivum* | Ensembl Plants | IWGSC |
| *Zea mays* | Ensembl Plants | B73_RefGen_v4 |
| *Zizania latifolia* | RiceRelativesGD | v1 |

**Supporting Table S4.** PBJelly gap filling summary statistics for the northern wild rice (NWR; *Zizania palustris*) *de novo* genome assembly.

| Scaffolds | Scaffolds with gaps | Scaffolds without gaps | Scaffolds with Ns | Scaffolds without Ns | Length of Gaps |
| --- | --- | --- | --- | --- | --- |
| Sequences | 2,183 | NA | 8,024 | NA | 5,841 |
| Minimum | 2 | 2 | 2 | 2 | 25 |
| 1<sup>st</sup> quartile | 4,605 | 4,605 | 21,086 | 21,086 | 100 |
| Median | 13,214 | 13,214 | 72,153 | 72,153 | 1,000 |
| Mean | 591,143 | 589,609 | 160,408 | 160,408 | 573.32 |
| 3<sup>rd</sup> quartile | 36,141 | 36,055 | 194,607 | 194,607 | 1,000 |
| Maximum | 118,081,501 | 117,893,876 | 3,907,081 | 3,907,081 | 1,000 |
| Total | 1,290,465,226 | 1,287,116,452 | 1,287,116,452 | 1,287,116,452 | 3,348,774 |
| N50 | 99,040,887 | 98,555,335 | 382,721 | 382,721 | 1,000 |
| N90 | 39,128,245 | 39,050,220 | 84,273 | 84,273 | 1,000 |
| N95 | 4,353,306 | 4,339,731 | 51,001 | 51,001 | 100 |

**Supporting Table S5.** Summary statistics for the northern wild rice (NWR; *Zizania palustris*) transcriptome assembly utilizing RNA-Seq data from eight different tissue types.

|  |  |
| --- | --- |
| Total # of transcripts | 689,344 |
| Total # of transcripts post-clustering | 624,117 |
| Number of “genes” | 418,924 |
| N50 contig length | 1,484 bp |
| Median contig length | 381 bp |
| Mean contig length | 783.38 bp |
| Total assembled sequence | 540,015,033 bp |

**Supporting Table S6.**
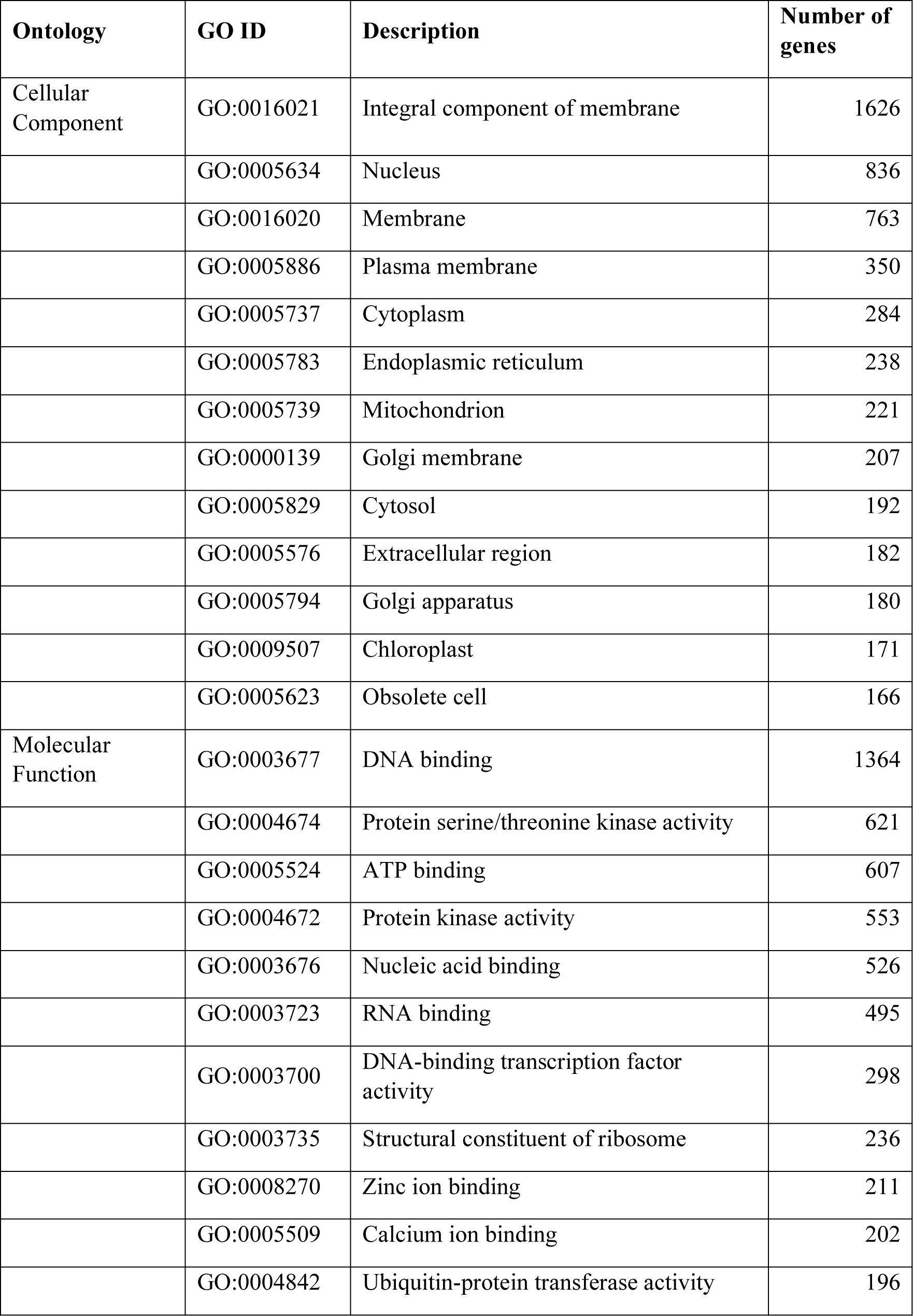
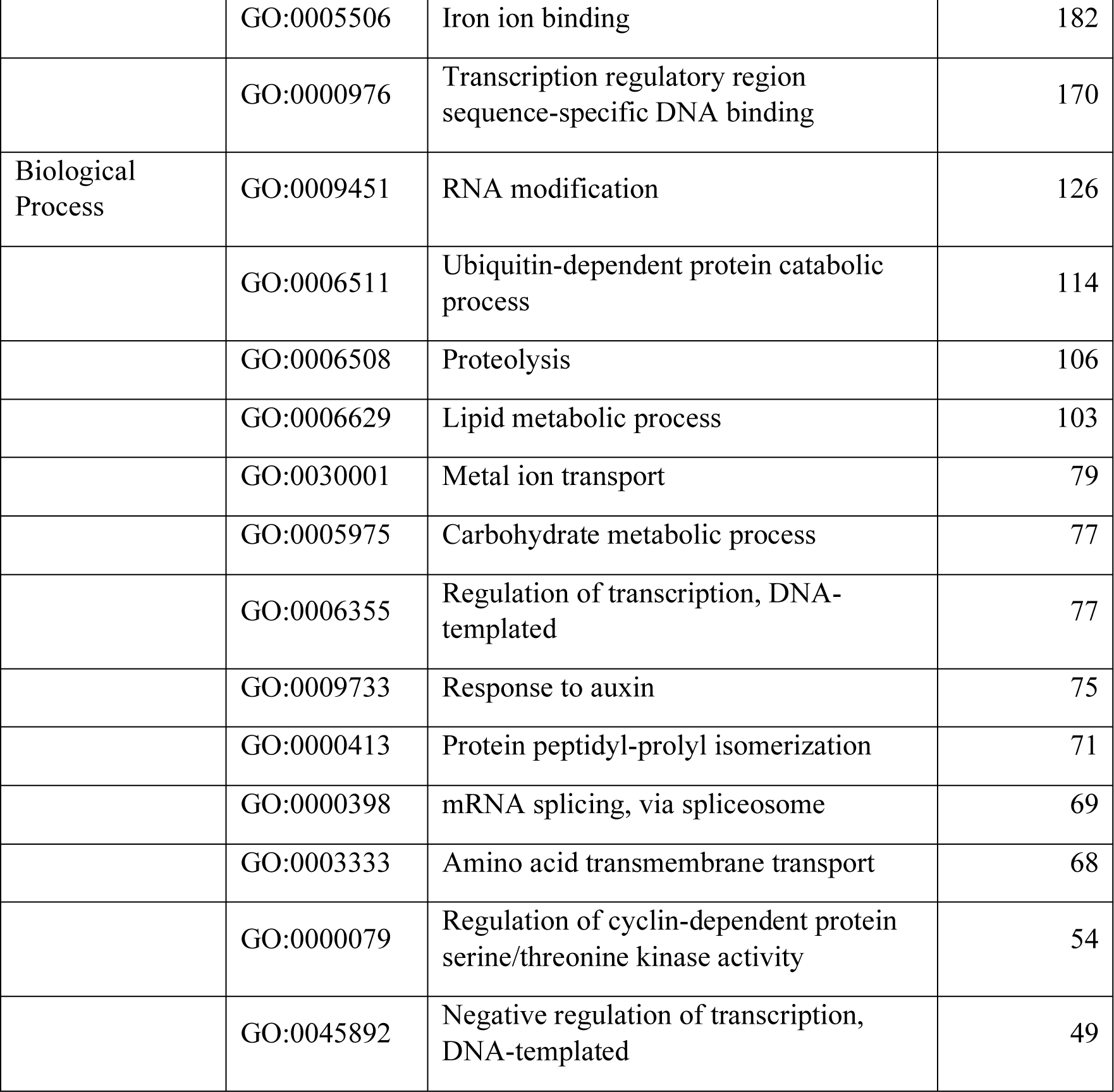
Major gene ontology (GO) terms for cellular component, molecular function, and biological process ontologies for the northern wild rice (NWR; *Zizania palustris*) genome annotation.

**Supporting Table S7:** Summary of the repeat element content in the northern wild rice (NWR; *Zizania palustris*) genome assembly as identified by RepeatMasker.

|  | Length occupied (bp) | % of sequence |
| --- | --- | --- |
| SINEs: | 97,530 | 0.01 % |
| ALUs | 0 | 0.00 % |
| MIRs | 0 | 0.00 % |
| LINEs: | 9,556,847 | 0.74 % |
| LINE1 | 8,224,354 | 0.64 % |
| LINE2 | 76,328 | 0.01 % |
| L3/CR1 | 262 | 0.00 % |
| LTR elements: | 763,436,744 | 59.24 % |
| ERVL | 90 | 0.00 % |
| ERVL-MaLRs | 0 | 0.00 % |
| ERV-classI | 21,1476 | 0.02 % |
| ERV-classII | 1,413 | 0.00 % |
| DNA elements: | 73,874,005 | 5.73 % |
| hAT-Charlie | 1,950 | 0.00 % |
| TcMar-Tigger | 33 | 0.00 % |
| Unclassified: | 137,342,785 | 10.66 % |
| Total interspersed repeats: | 984,307,911 | 76.38 % |
| Small RNA: | 66,477 | 0.01 % |
| Satellites: | 19,101 | 0.00 % |
| Simple repeats: | 5,296,521 | 0.41 % |
| Low complexity: | 755,600 | 0.06 % |

1-SINE: Small Interspersed Nuclear Element; LINE: Long Interspersed Nuclear Element; LTR=Long Terminal Repeat.

**Supporting Table S8.**
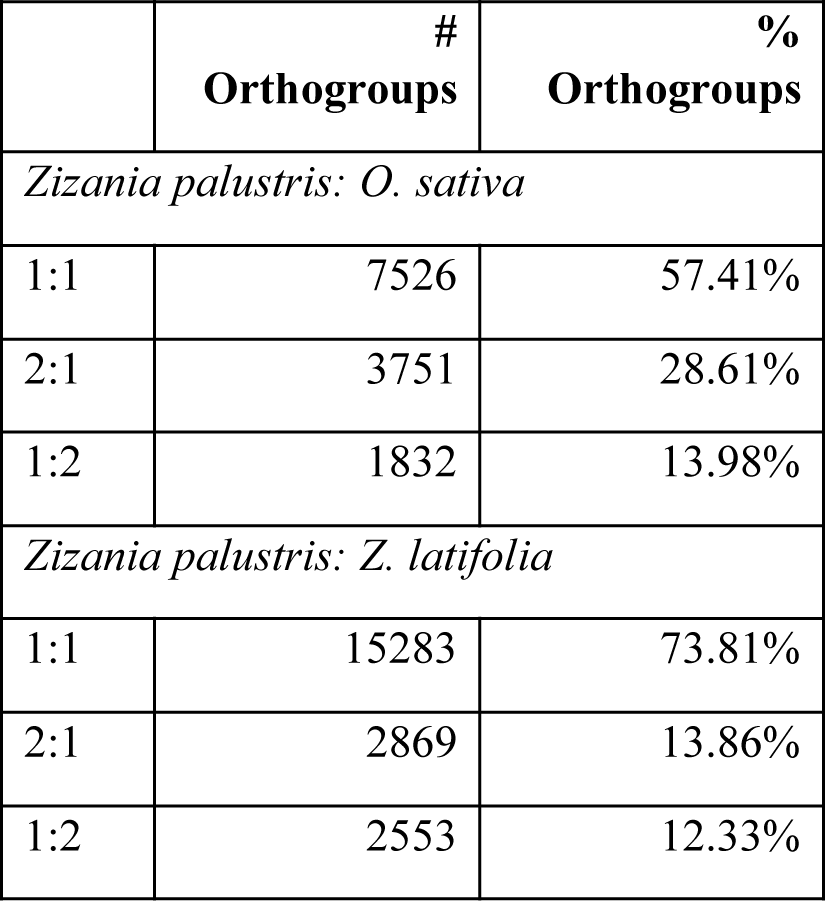
Distribution of 1:1, 1:2, and 2:1 orthogroup relationships between northern wild rice (NWR; *Z. palustris*), *O. sativa*, and *Z. latifolia*.

**Supporting Table S9.** Summary of syntenic blocks detected through the comparison of the northern wild rice (NWR; *Zizania palustris*) genome with rice (*Oryza sativa*), *Zizania latifolia*, and itself.

| Genome comparisons | No. of syntenic blocks | No. of gene pairs/block | No. of syntenic gene pairs | Mean block length (1000 nt) |
| --- | --- | --- | --- | --- |
| NWR vs. rice | 262 | 135.22 | 35,430 | 6,416.19 vs. 3,125.41 |
| NWR vs. *Z. latifolia* | 1,321 | 14.60 | 19,293 | 6,440.39 vs. 200.38 |
| NWR vs. NWR | 118 | 138.77 | 16,375 | 8,579.62 |

**Supporting Table 10.**
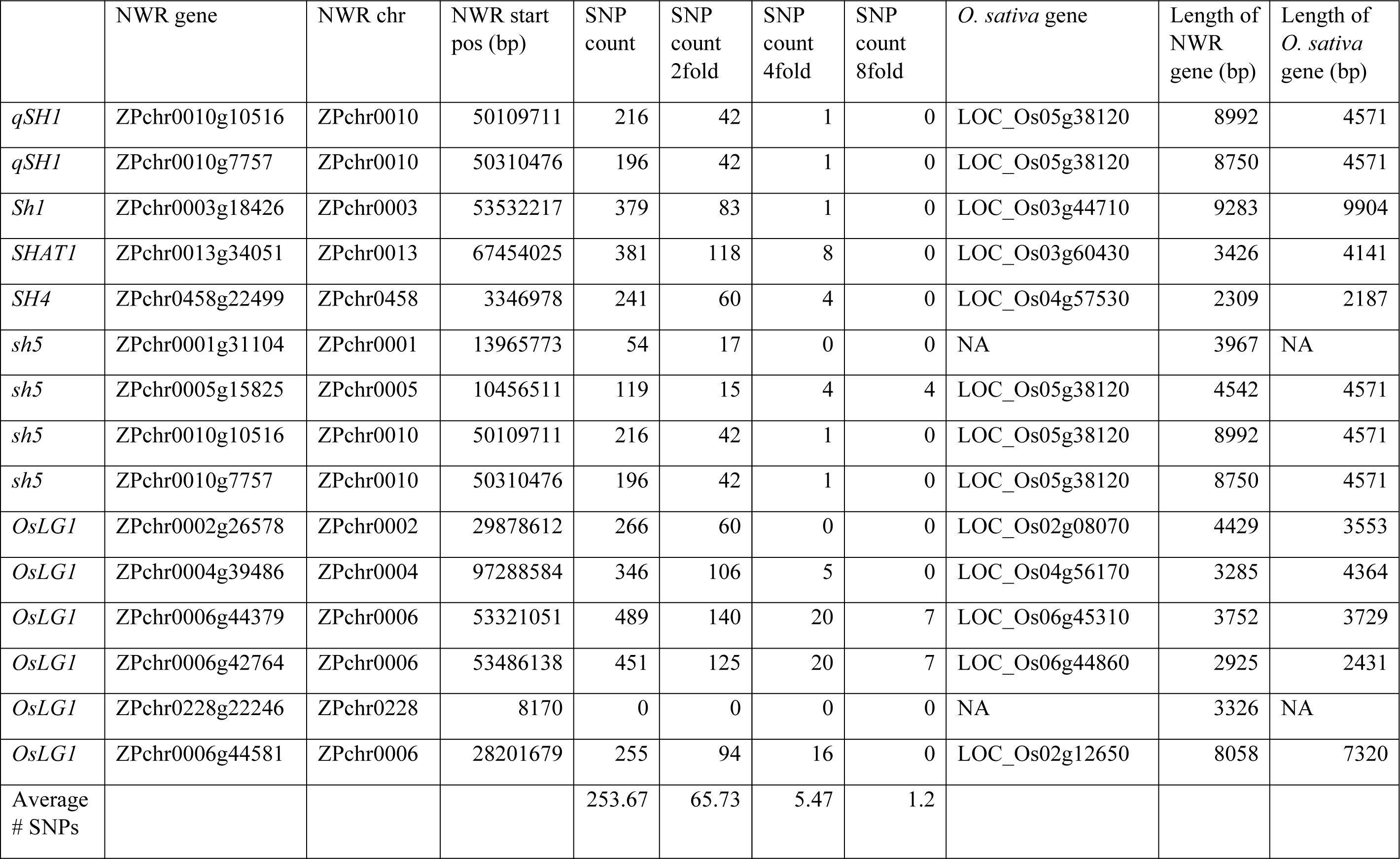
A comparison of fifteen putative NWR shattering genes and their orthologs in *O. sativa.* The number of SNPs in a 2 Mb window (1 Mb up- and downstream from the start position of each gene). The SNPs were identified using GBS data from Shao *et al*. (2020). The selected NWR genes were chosen based on their inclusion in Table 3.

|  | NWR gene | NWR chr | NWR start pos (bp) | SNP count | SNP count 2fold | SNP count 4fold | SNP count 8fold | *O. sativa* gene | Length of NWR gene (bp) | Length of *O. sativa* gene (bp) |
| --- | --- | --- | --- | --- | --- | --- | --- | --- | --- | --- |
| *qSH1* | ZPchr0010g10516 | ZPchr0010 | 50109711 | 216 | 42 | 1 | 0 | LOC_Os05g38120 | 8992 | 4571 |
| *qSH1* | ZPchr0010g7757 | ZPchr0010 | 50310476 | 196 | 42 | 1 | 0 | LOC_Os05g38120 | 8750 | 4571 |
| *Sh1* | ZPchr0003g18426 | ZPchr0003 | 53532217 | 379 | 83 | 1 | 0 | LOC_Os03g44710 | 9283 | 9904 |
| *SHAT1* | ZPchr0013g34051 | ZPchr0013 | 67454025 | 381 | 118 | 8 | 0 | LOC_Os03g60430 | 3426 | 4141 |
| *SH4* | ZPchr0458g22499 | ZPchr0458 | 3346978 | 241 | 60 | 4 | 0 | LOC_Os04g57530 | 2309 | 2187 |
| *sh5* | ZPchr0001g31104 | ZPchr0001 | 13965773 | 54 | 17 | 0 | 0 | NA | 3967 | NA |
| *sh5* | ZPchr0005g15825 | ZPchr0005 | 10456511 | 119 | 15 | 4 | 4 | LOC_Os05g38120 | 4542 | 4571 |
| *sh5* | ZPchr0010g10516 | ZPchr0010 | 50109711 | 216 | 42 | 1 | 0 | LOC_Os05g38120 | 8992 | 4571 |
| *sh5* | ZPchr0010g7757 | ZPchr0010 | 50310476 | 196 | 42 | 1 | 0 | LOC_Os05g38120 | 8750 | 4571 |
| *OsLG1* | ZPchr0002g26578 | ZPchr0002 | 29878612 | 266 | 60 | 0 | 0 | LOC_Os02g08070 | 4429 | 3553 |
| *OsLG1* | ZPchr0004g39486 | ZPchr0004 | 97288584 | 346 | 106 | 5 | 0 | LOC_Os04g56170 | 3285 | 4364 |
| *OsLG1* | ZPchr0006g44379 | ZPchr0006 | 53321051 | 489 | 140 | 20 | 7 | LOC_Os06g45310 | 3752 | 3729 |
| *OsLG1* | ZPchr0006g42764 | ZPchr0006 | 53486138 | 451 | 125 | 20 | 7 | LOC_Os06g44860 | 2925 | 2431 |
| *OsLG1* | ZPchr0228g22246 | ZPchr0228 | 8170 | 0 | 0 | 0 | 0 | NA | 3326 | NA |
| *OsLG1* | ZPchr0006g44581 | ZPchr0006 | 28201679 | 255 | 94 | 16 | 0 | LOC_Os02g12650 | 8058 | 7320 |
| Average # SNPs |  |  |  | 253.67 | 65.73 | 5.47 | 1.2 |  |  |  |

**Supporting Figure S1.**
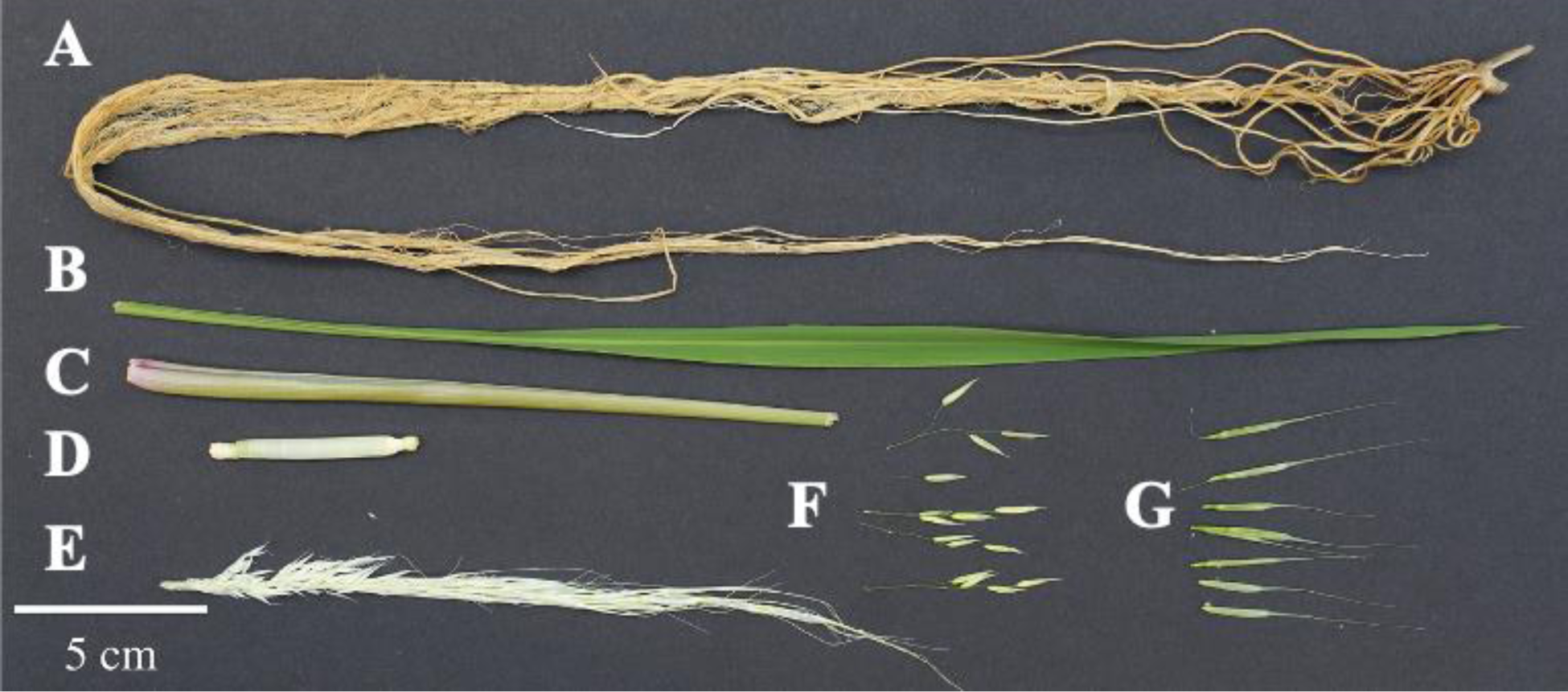
Examples of northern wild rice (NWR: *Zizania palustris*) tissues, including: A. root, B. leaf, C. leaf sheath, D. stem, E. whole un-emerged panicle, F. male florets, and G. seed, which were harvested for sequencing (RNA-seq).

**Supporting Figure S2.**
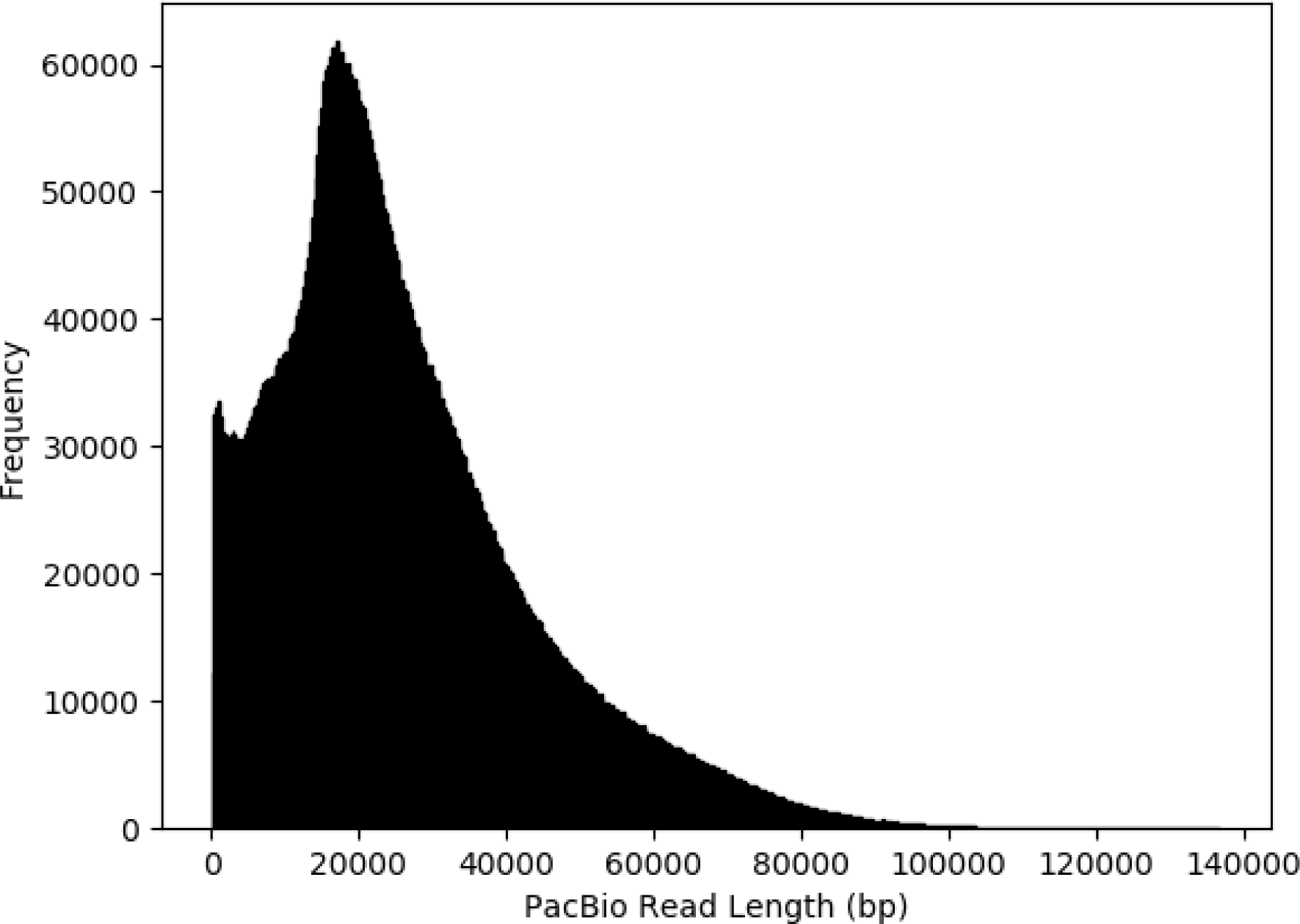
Distribution of PacBio sequencing read lengths of northern wild rice (NWR: *Zizania palustris*) cultivar, Itasca-C12.

**Supporting Figure 3.**
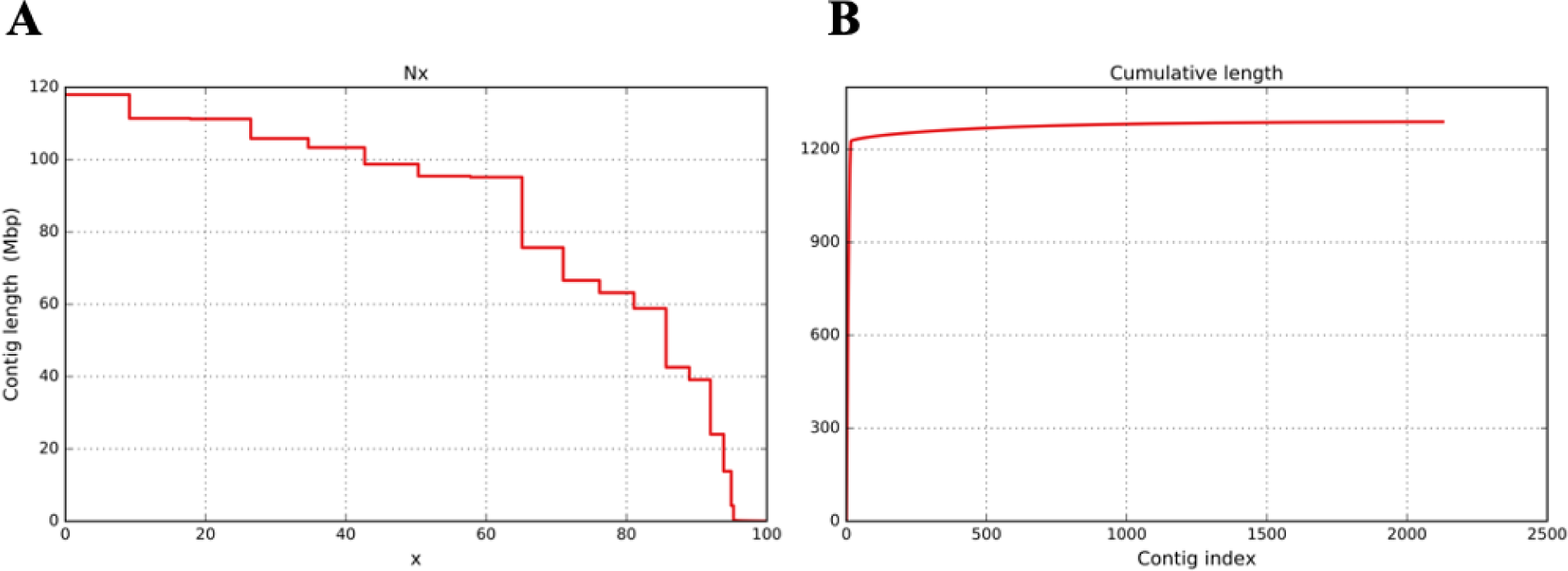
Northern wild rice (NWR; *Zizania palustris*) genome assembly statistics. A. Nx plot showing the percentage of the genome assembly covered by each scaffold’s length in Mb, where scaffolds are ordered. B. Plot showing the contributions of the 2,183 scaffolds to the overall genome assembly size.

**Supporting Figure S4.**
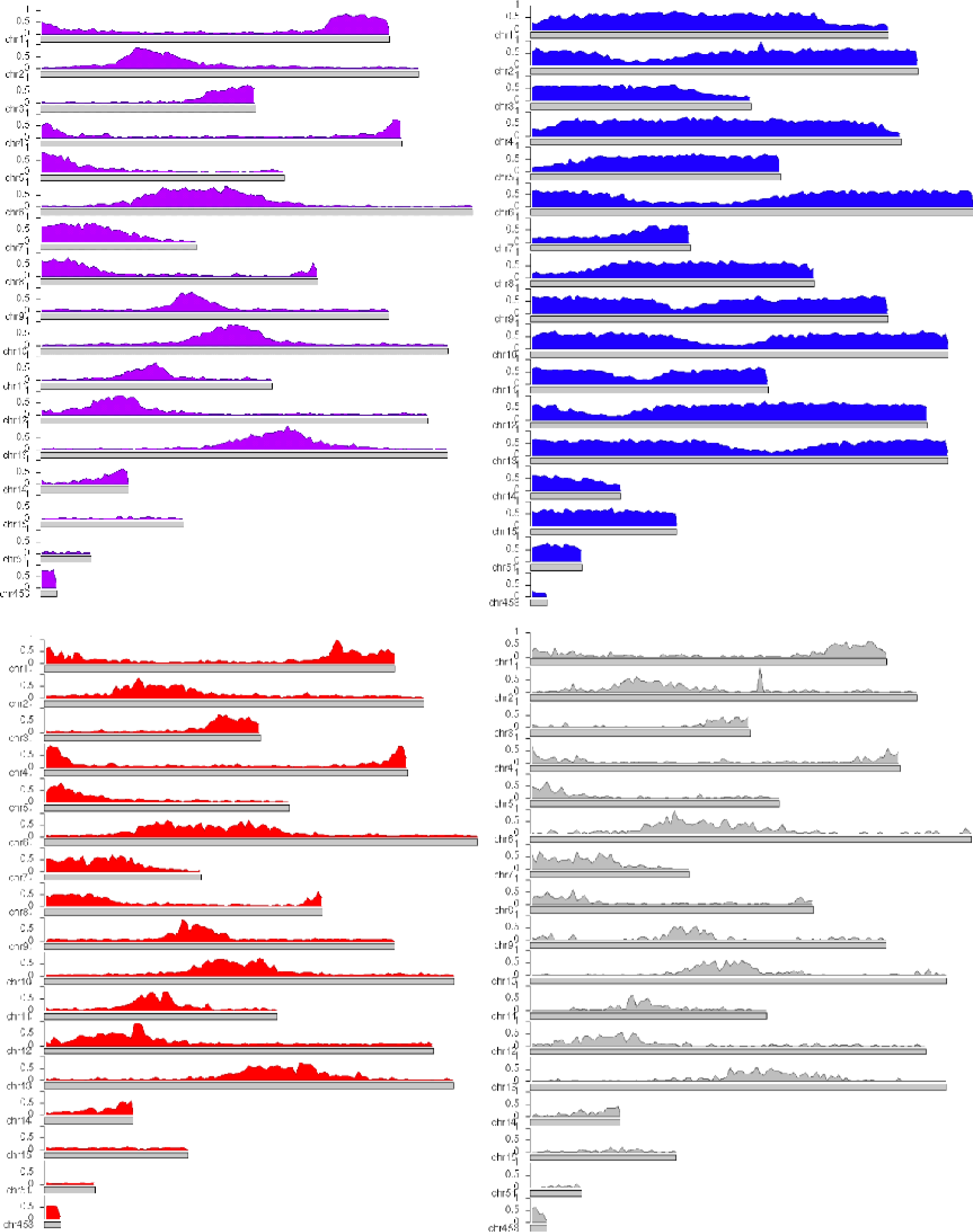
Density plots representing the distribution and density of northern wild rice 1034 (NWR; *Zizania palustris*) A. predicted genes; B. long-terminal repeats (LTR); C. DNA elements; and D. long-interspersed nuclear elements (LINES).

**Supporting Figure S5.**
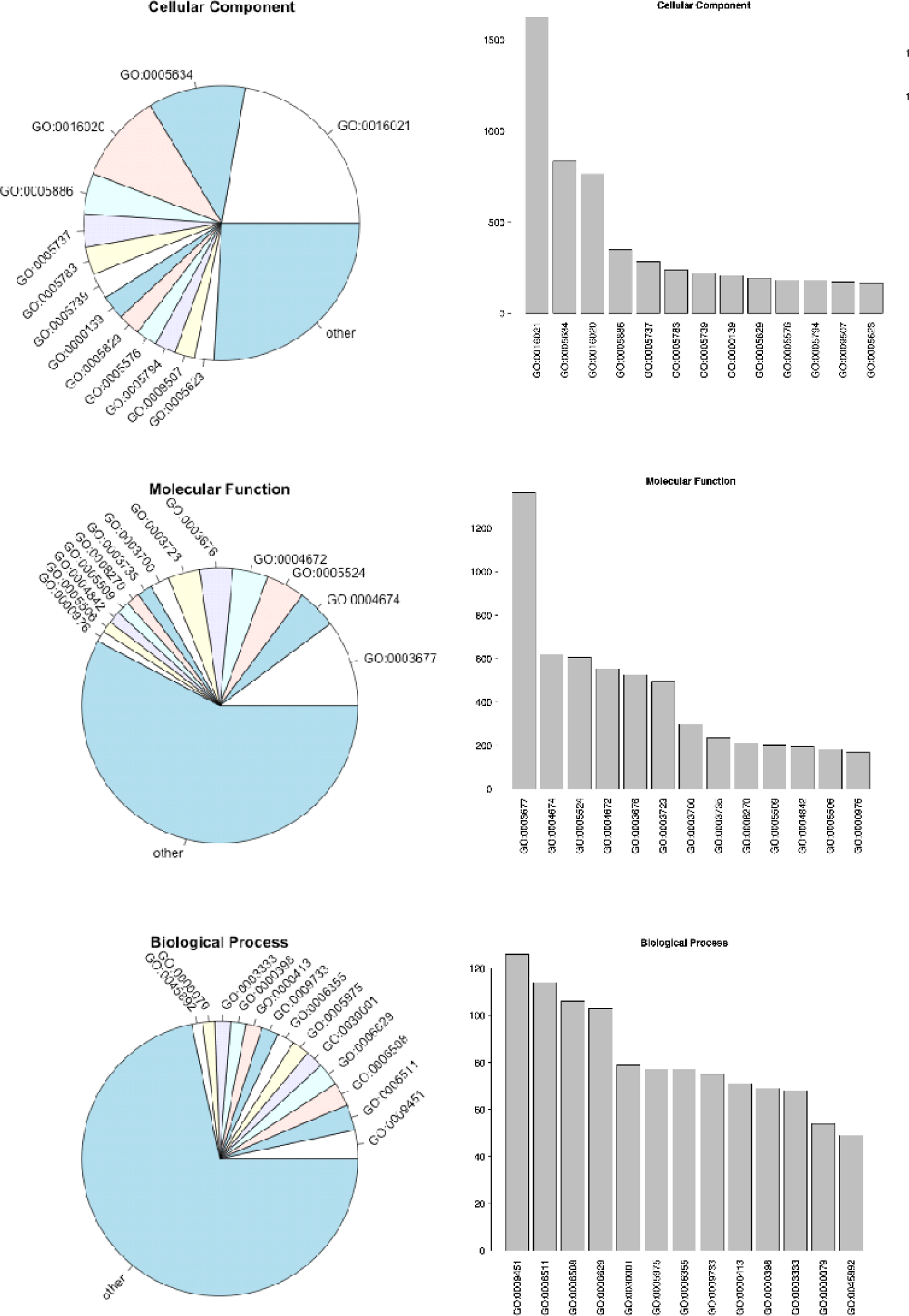
Composition of northern wild rice (NWR; *Zizania palustris*) gene function based on gene ontology (GO) terms. Distributions are shown for A. Cellular Component (CC), B. Molecular 1039 Function (MF), and C. Biological Process (BP) ontologies.

**Supporting Figure S6.**
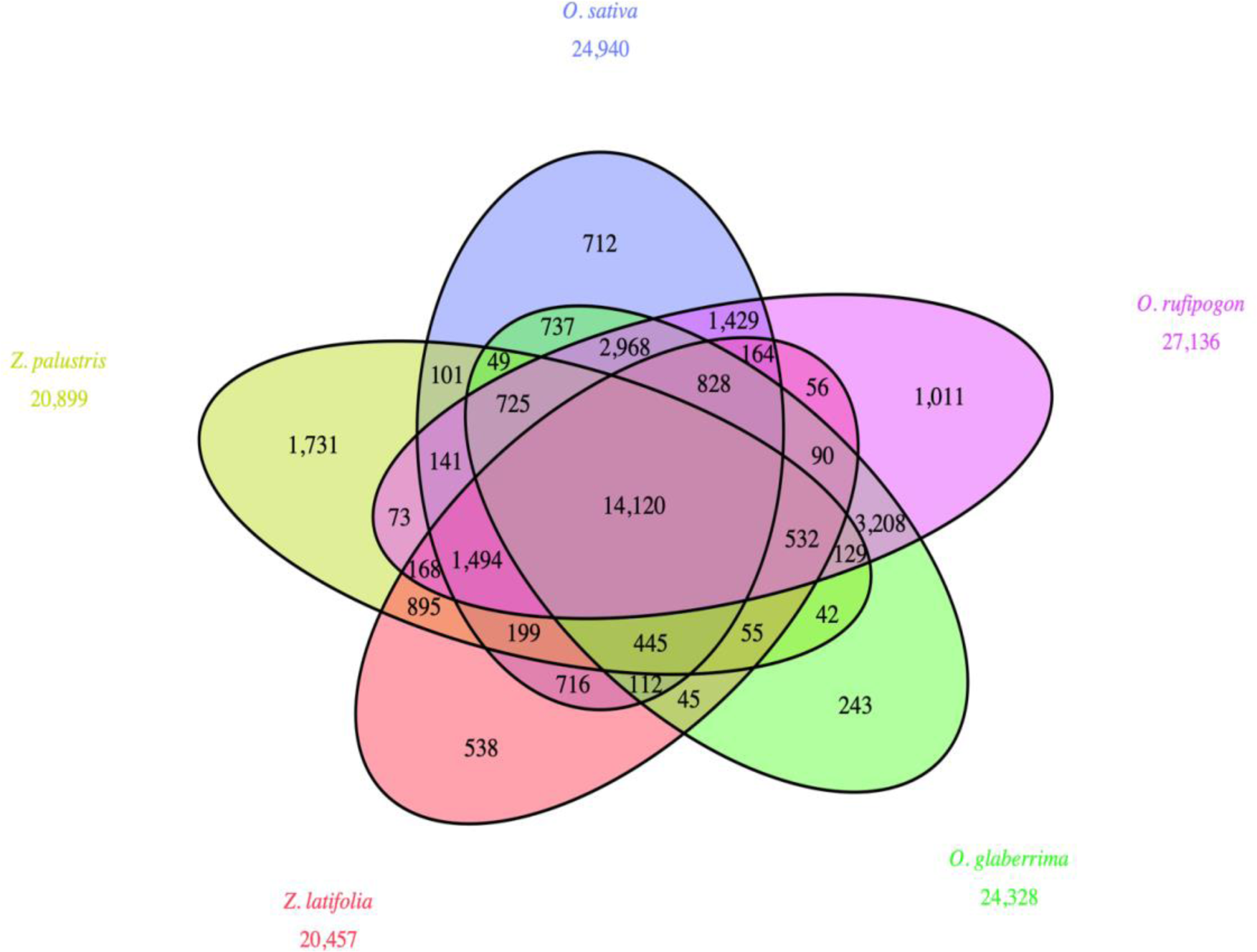
Venn diagram showing the number of orthogroups for *O. sativa*, *O. rufipogon*, *O.* 1044 *glaberrima*, northern wild rice (NWR; *Zizania palustris*), and *Z. latifolia*.

**Supporting Figure S7.**
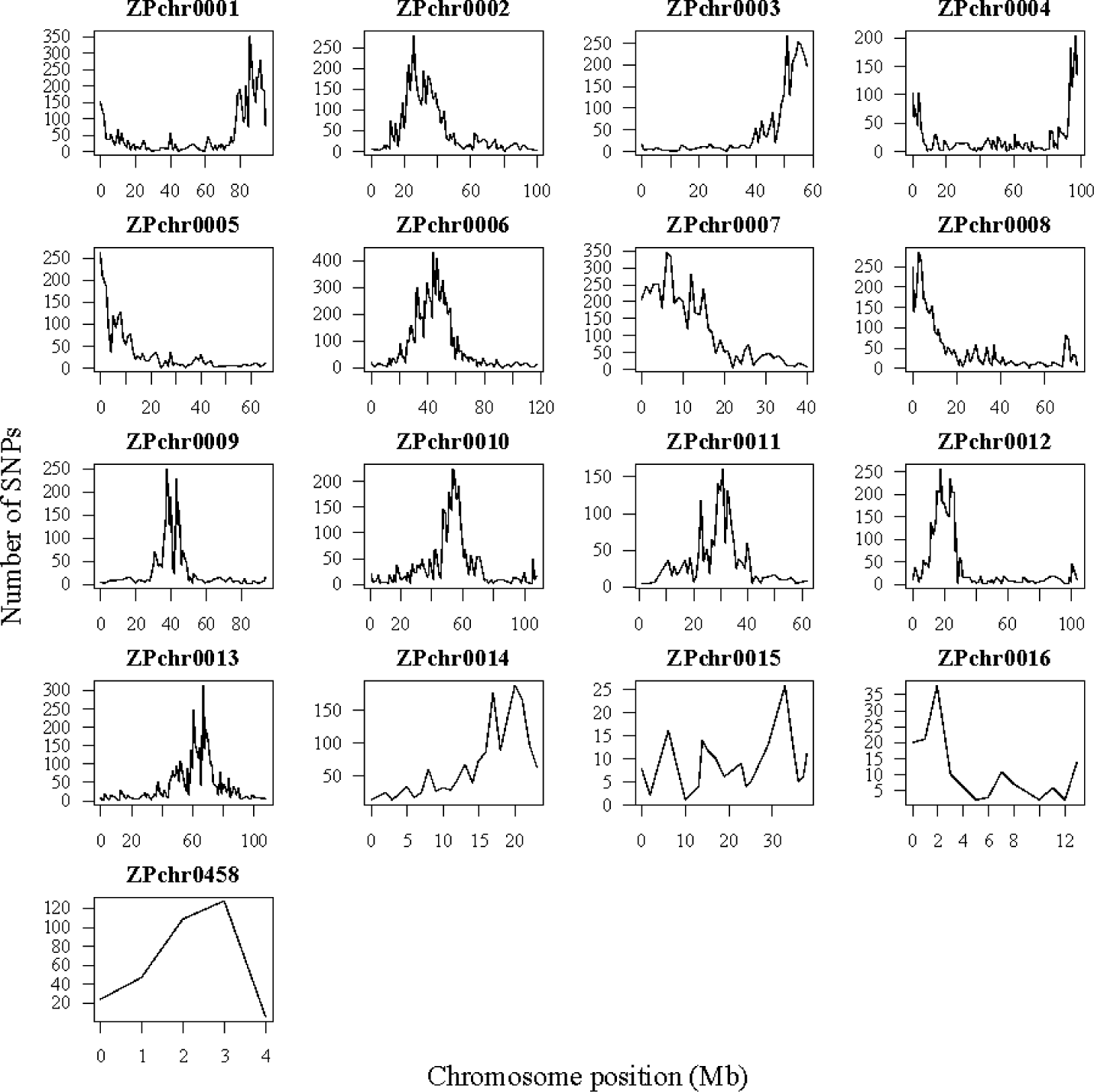
The distribution of SNPs along the seventeen major NWR chromosomes in 1Mb come from Shao *et al*. (2020) and were not downsampled (e.g., represent the original depth of ample for 8 total samples).

## Notes

### Competing Interest Statement

The authors have declared no competing interest.

